# Circulating miR-181 is a prognostic biomarker for amyotrophic lateral sclerosis

**DOI:** 10.1101/833079

**Authors:** Iddo Magen, Nancy Sarah Yacovzada, Eran Yanowski, Anna Coenen-Stass, Julian Grosskreutz, Ching-Hua Lu, Linda Greensmith, Andrea Malaspina, Pietro Fratta, Eran Hornstein

## Abstract

Amyotrophic lateral sclerosis (ALS) is a relentless neurodegenerative syndrome of the human motor neuron system, for which no effective treatment exists. Variability in the rate of disease progression limits the efficacy of ALS clinical trials, suggesting that developing of better biomarkers for prognosis will facilitate therapeutic progress. Here, we applied unbiased next-generation sequencing to investigate the potential of plasma cell-free microRNAs as biomarkers of ALS prognosis, in 252 patients with detailed clinical-phenotyping. First, we identified miRNAs, whose plasma levels remain stable over the course of disease in a longitudinal cohort of 22 patients. Next, we demonstrated that high levels of miR-181, a miRNA enriched in neurons of the brain and spinal cord, predicts a >2 fold risk of death in discovery cohort (126 patients) and an independent replication cohort (additional 122 patients). miR-181 performance is comparable with the established neurofilament light chain (NfL) biomarker and when combined together, miR-181+NfL establish a novel RNA-protein biomarker pair with superior prediction capacity of ALS prognosis. Therefore, plasma miR-181 predicts ALS disease course, and a novel miRNA-protein biomarker approach, based on miR-181+NfL, boosts precision of patient stratification and may greatly enhance the power of clinical trials.

**One Sentence Summary:** plasma miR-181 levels indicate high mortality risk in ALS patients.

## Introduction

Amyotrophic lateral sclerosis (ALS) is a devastating neurodegenerative disorder of the motor neuron system, for which no effective disease-modifying treatment exists. ALS is characterized by a significant variability in progression rates ^1, 2^, posing a significant challenge for patient stratification in clinical trials. Thus, reliable predictors of disease progression would be invaluable for ALS patient stratification prior to enrolment in clinical trials. Ideal biomarkers should remain stable during the course of disease, be detectable in accessible tissue, and also be easily measurable. To date, intensive research has identified only a few potential blood-based ALS biomarkers ^3–5^, including cell-free neurofilaments ^6–8^, and pro-inflammatory cytokines ^9–11^. Neurofilament light chain (NfL) was the first blood biomarker to aid in predicting ALS progression rate, but further markers are needed to improve stratification and allow for more effective trials. microRNAs (miRNAs) are endogenous non-coding RNAs that are essential for motor neuron survival and have been shown to be globally downregulated in post mortem ALS motor neurons ^12–14^. While circulating miRNA profiles have been previously characterized in ALS ^15–19^, the potential of miRNA biomarkers for ALS prognosis, and as readout of disease progression has not been fully explored.

Here, we take a hypothesis-free approach by applying next generation sequencing (RNA-seq) to comprehensively study plasma miRNAs in a large cohort of 252 ALS cases. These studies focused our attention on the miR-181 family, which are expressed from two homologs, polycistronic genes, *mir-181a-1/b-1* (human chromosome 9) and *mir-181a-2/b-2* (human chromosome 1). The mature miR-181 species are functionally identical in silencing a single set of mRNA targets. We reveal that miR-181 levels predict disease progression in large discovery and a replication cohort, and demonstrate the effectiveness of combining miR-181 with established neurofilament light chain as a prognostic biomarker pair for ALS.

## Results

### Longitudinal study of circulating miRNAs in ALS

In this work, we sought to explore blood-borne miRNAs as potential prognostic biomarkers for ALS. We used unbiased next generation sequencing to investigate, without an *a priori* bias, the comprehensive landscape of plasma miRNAs in 252 ALS patients, for which documented clinical and demographic information is available (Table 1).

**Table 1.**
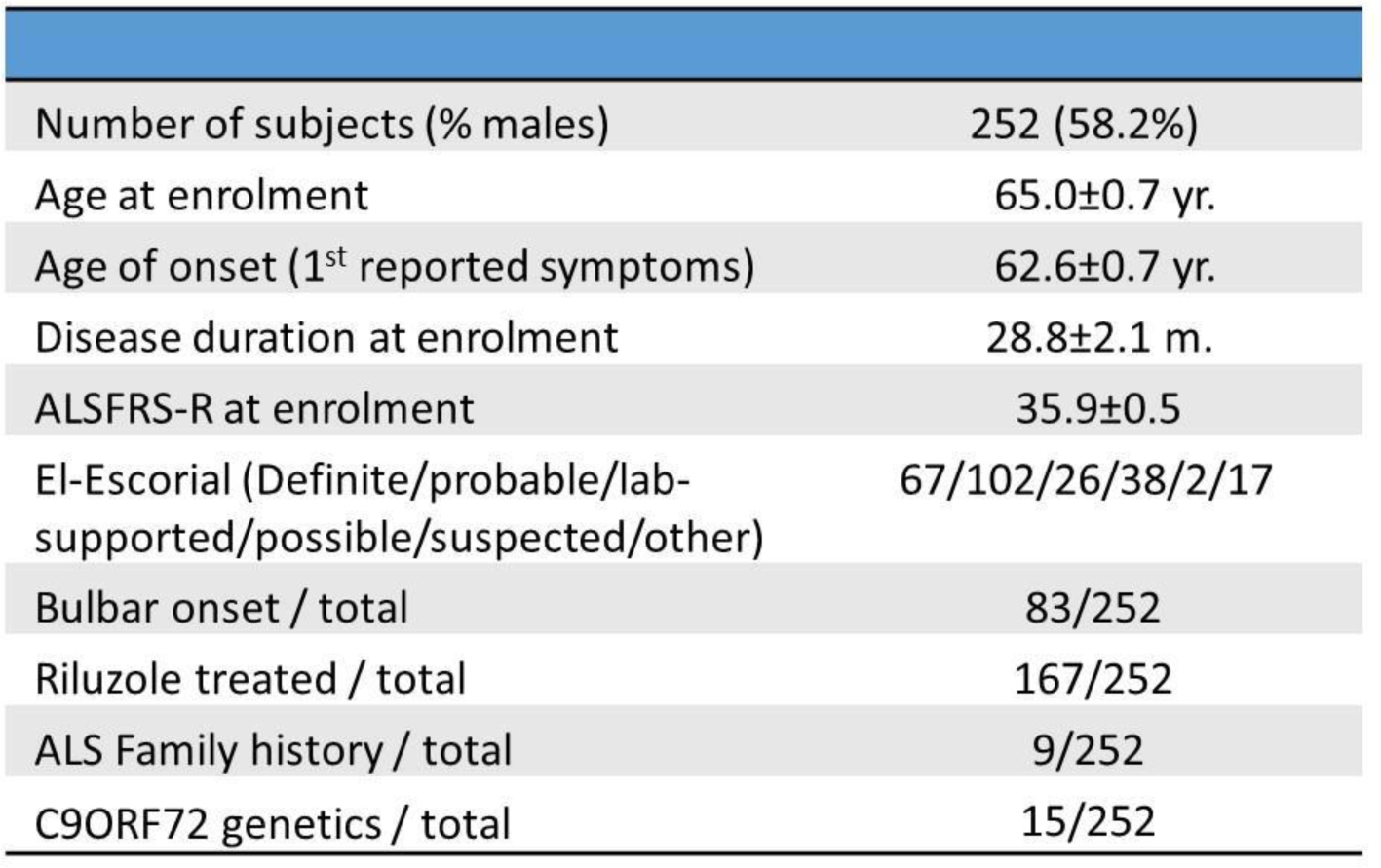
Summary of demographic and clinical characteristics of ALS samples used for the survival study. ALSFRS-R: ALS functional rating scale. Data are presented as mean ± SEM.

A crucial feature for a prognostic biomarker is its stability across the disease course. We therefore initially investigated a longitudinal sample cohort of 22 patients (clinical data in Table 2), with four longitudinal blood samples taken (t_1_-t_4_) during the course of 30 months (2.5 years). 88 samples (corresponding to the first cohort of 22 patients), were prepared from total plasma RNA, as previously described ^20^, and profiled by RNA-seq for miRNA levels. Linear miRNA quantification was achieved via unique 12-nucleotide molecular identifiers (UMIs). miRNAs with ≥50 UMIs in at least 60% of the samples (>53 out of 88 samples) were considered above noise level. Thus, of 2008 miRNAs aligned to the human genome (GRCh37/hg19), 187 passed the threshold we set (see Table S1). To reduce noisy miRNAs, we next excluded from further analysis 58 miRNAs with high variability (t4/t1 standard error ratio ≥ 0.2, Figure 1A, y-axis). For example, miR-181a-5p variability across individual patients is limited, relative to that of miR-1-3p (F test for variance = 20.9, p<0.0001, Fig. 1B). We identified 125 miRNA candidate biomarkers, whose plasma levels were relatively stable during disease progression in the same sub-cohort of 22 longitudinal samples (Figure 1A, Table S1) that could be tested as candidate prognostic biomarkers. In addition, four miRNAs, whose levels increased during the course of disease, were subjected validation in a separated replication longitudinal cohort (N=26 patients) and may serve measures of functional decline over the course of the disease (Figures S1, S2, S3; Table S3).

**Figure 1.**
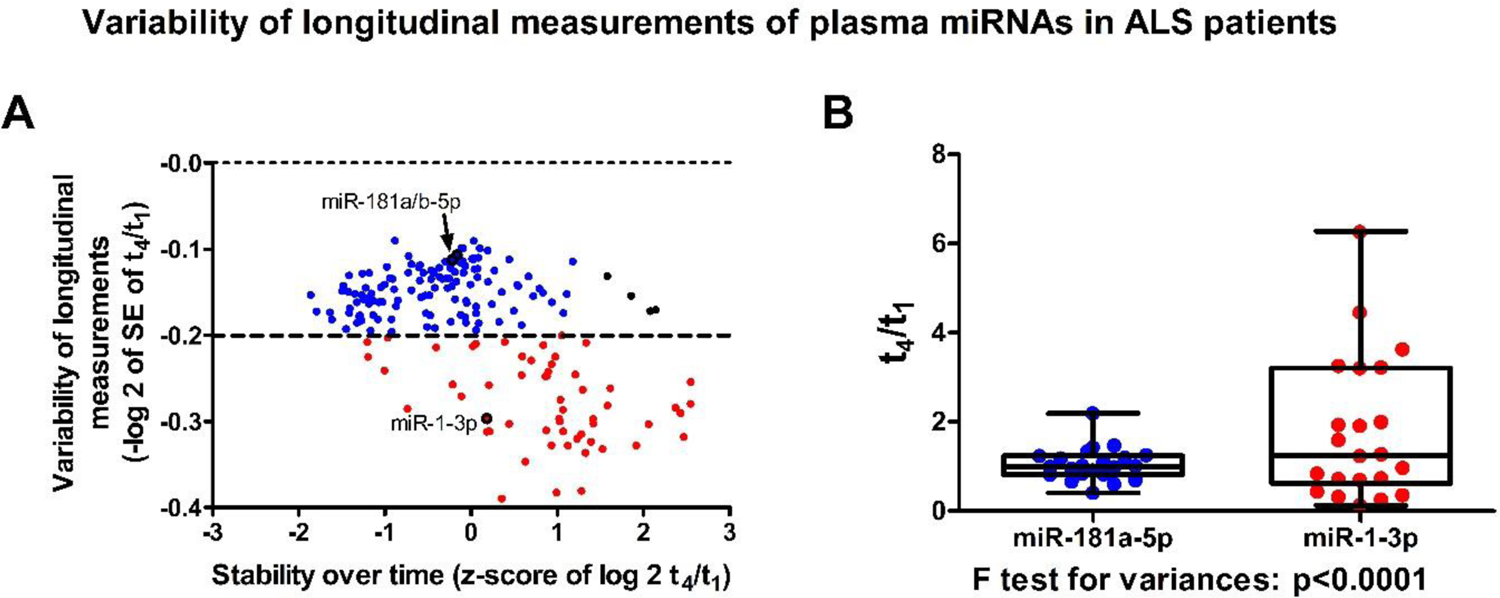
Assessment of plasma miRNA stability during ALS course. (A) The x-axis denotes the standardized change in miRNA levels between the first and last measurements (number of standard deviations (SD) for log 2-transformed t_4_/t_1_ ratios), relative to the average change of all 187 sequenced miRNAs. The y-axis denotes the variability in measurements, per-miRNA for 187 sequenced species between the 22 individuals (-log 2-transformed values of the standard error of t_4_/t_1_ ratios). Features above (blue) or below (red) the stability threshold set at −0.2 units. **(B)** Variability of miR-181a-5p and miR-1-3p between fourth and first phlebotomy (t4/t1) in 22 ALS patients. F test p<0.0001.

**Table 2.** Summary of demographic and clinical characteristics of ALS samples used for the longitudinal study. ALSFRS-R: ALS functional rating scale. Data are presented as mean ± SEM.

|  | Cohort I | Cohort II |
| --- | --- | --- |
| Number of subjects (% males) | 22 (81.8%) | 26 (53.8%) |
| Age at enrolment | 65.0±1.5 yr. | 63.5±2.4 yr. |
| Age of onset (1<sup>st</sup> reported symptoms) | 62.0±1.7 yr. | 60.4±2.4 yr. |
| Disease duration at enrolment | 37.2±8.2m | 37.0±8.5 m. |
| ALSFRS-R at enrolment | 41.0±1.2 | 36.6±1.4 |
| El-Escorial (Definite/probable/lab-supported/possible/other) | 4/7/6/5/0 | 3/14/3/3/3 |
| Bulbar onset / total | 8/22 | 6/26 |
| Riluzole treated / total | 13/22 | 15/26 |
| ALS Family history / total | 1/22 | 1/26 |
| C9ORF72 genetics / total | 0/22 | 3/26 |

### Discovery of circulating miRNAs as potential biomarkers for ALS prognosis

For the main interest of the current study in prognosis analysis, we focused on the 125 miRNAs that displayed stable plasma levels over time. These 125 miRNAs were further investigated in a cohort of 252 patients, for which a single blood sample was collected at enrolment. We randomly split the cohort into two sub-cohorts of 126 patients each, with comparable demographic and clinical features (Figure S4).

We performed next generation sequencing on the first cohort of 126 patients termed “discovery cohort”, holding out an equally-sized “replication cohort” for validation. Out of the 125 candidate miRNAs, we excluded 19 miRNAs, which did not pass the minimal UMI threshold or QC (Figure S5). Optimal cut-off values were determined for 106 miRNAs predictors, for dichotomizing continuous expression levels to binary (high/low), by iterative testing of the capacity to predict patient survival (time elapsed to death, using *Evaluate Cutpoints* algorithm ^21^). 19 additional miRNA were excluded at the QC step (methods). Nine of the remaining 87 miRNAs predicted prognosis in a significant manner, when survival was calculated from either onset (defined as first documented symptoms) or enrollment (Figure 2A, B; Table S1). We further tested the prediction capacity of combinations of miRNAs considering this way potential cooperative information in evaluation of all 36 miRNA pairs 20 out of 36 miRNA pairs predicted prognosis comparably or superior to individual miRNAs (logrank p value ≤0.01, Figure 2A, B, S6, Table S1).

**Figure 2.**
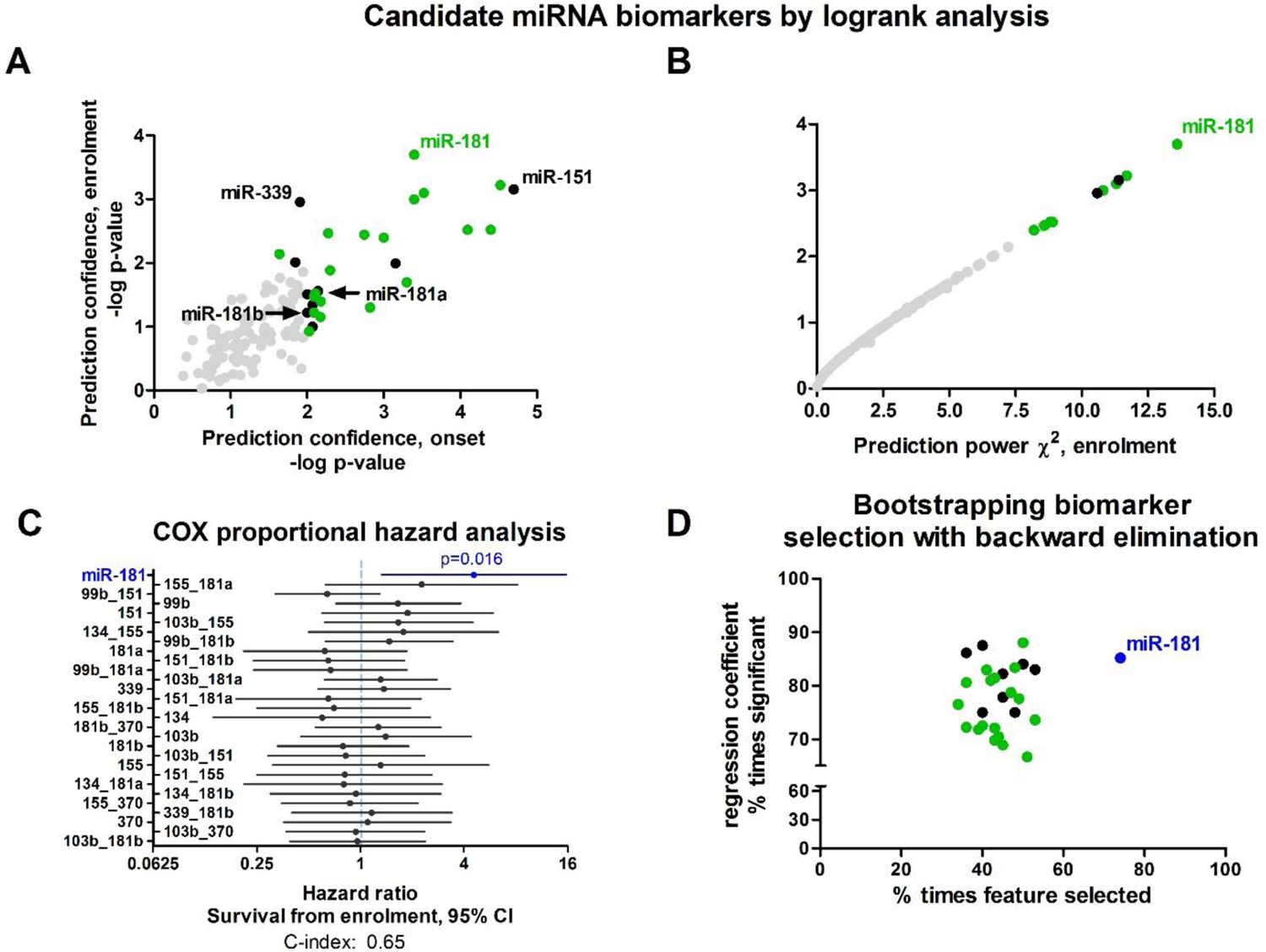
Identification of candidate miRNAs that predict ALS patient survival. (A) Scatter plot, assessing separation of survival by high and low (dichotomized) levels of 123 miRNA features (i.e., single miRNAs and miRNA pairs). Logrank test from study enrollment (y-axis) or first symptoms (onset, x-axis). Log 10 transformed p-values of logrank chi^2 values. The optimal threshold was calculated per miRNA in a discovery cohort of 126 patients by Evaluate Cutpoints algorithm ^21^. Single miRNA (black, namely, miR-103b-3p, miR-134-5p, miR-151-5p, miR-155-5p, miR-181a-5p, miR-181b-5p, miR-339-3p, miR-370-3p, miR-99b-5p) or miRNA pairs (green), displaying a p-value ≤0.01 (log 10 transformed values ≥ 2), and grey: insignificant. The paired feature composed of miR-181a-5p with miR-181b-5p is called for simplicity miR-181 throughout the manuscript. (**B**) Scatter plot of effect (logrank chi^2 values, x-axis), against confidence (p-value, y-axis). Color code as in panel A. (**C**) Forest plot of mortality hazard ratios calculated by multivariate COX study from enrolment for all significant miRNAs and miRNA pairs in Figure 2A (colored black and green). Blue features - displaying a p-value ≤0.05, black: insignificant by Wald test. (**D**) Bootstrap-based model selection according to Akaike’s information criteria (AIC). Backwards feature elimination for 29 features (miRNA or miRNA pairs) passing p-value ≤0.01 filtering in the logrank test (Figure 2A). Plot of the percentage of times each feature was selected (x-axis), against the percentage of times the Cox regression coefficient of this feature was significant in repeated measurements (y-axis).

The monthly mortality hazard ratio (HR) was calculated for 9 miRNAs and 20 miRNA-pairs in a multivariate Cox regression analysis, stratified by the disease stage (at enrollment) and age at onset (methods). This analysis allows calculation of an independent hazard ratio for each covariate (i.e., single miRNA or miRNA pair), while holding the other covariates constant. We report a risk of dying that is almost five times higher with high plasma levels of miR-181 (featuring two sister miRNAs, miR-181a-5p and miR-181b-5p, Figure 2B; hazard ratio (HR) = 4.55, 95% CI: 1.33 - 15.6, p = 0.016, Figure 2C). None of the other features reached a statistically significant signal. Noteworthy, assessment of miR-181 levels as a continuous variable, opposed to categorical one, did not contribute to prediction of mortality hazard.

Stepwise feature selection using bootstrap resampling procedure ^22^ is a rigorous scheme for the selection of robust survival outcome predictors, that has been used in ALS biomarker research ^23^. We therefore orthogonally selected candidate predictors using backward feature elimination, according to Akaike’s information criteria (AIC) across 100 bootstrap samples (Figure 2D; Table S1). miR-181 was the only feature satisfying bootstrap criteria (selected >70%, significant >85%). Taken together, these data identify miR-181 as the best miRNA predictor of survival in ALS patients by both traditional statistics (logrank analysis (Fig. 2A,B), multivariate Cox proportional hazard (Fig 2C) and by bootstrap model selection (Fig 2D).

### Validation of circulating miR-181 as biomarker for ALS prognosis

We next tested the capacity of miR-181 to separate survival curves of patients. Kaplan Meier curves revealed clear separation of survival between with high vs low miR-181 subgroups, based on plasma miR-181 levels at enrolment (discovery cohort: logrank chi^2 = 13.6, p=0.0002, Figure 3A; Table S2). The median patient survival associated with low miR-181 was 18.6 months, compared to 9 months associated with higher miR-181 levels. Thus, plasma miR-181 levels predict a substantial median survival difference of 9.6 months that is equivalent to a 207% increase in survival length for patients with lower plasma miR-181 levels. Comparable results were obtained when survival length was calculated from disease onset (Figure 3B).

**Figure 3.**
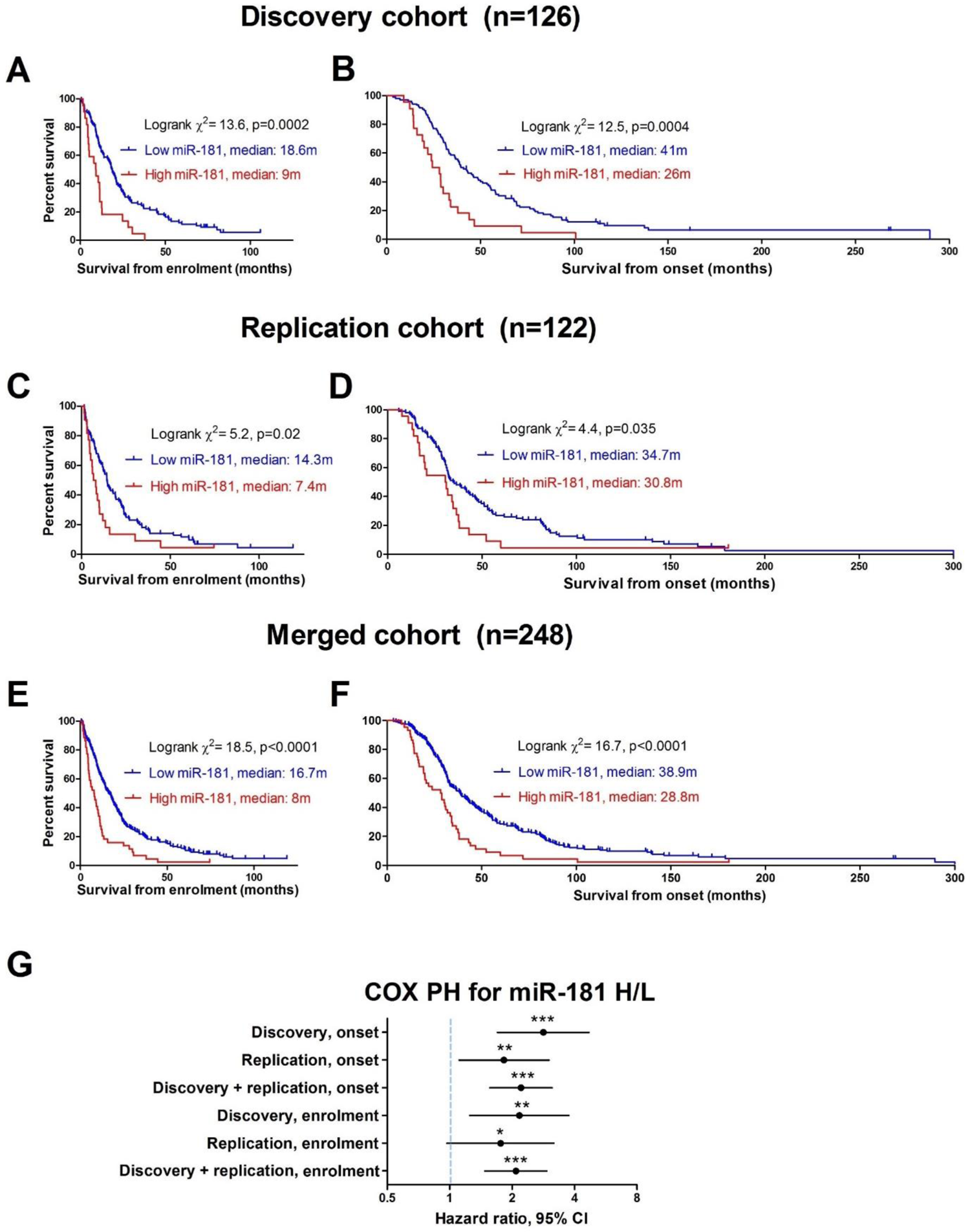
miR-181 is a prognostic biomarker of ALS. Cumulative survival (Kaplan-Meier) curves for miR-181 (104 patients with subthreshold (blue) vs. 22 patients with suprathreshold (red) miR-181 levels) in discovery cohort from enrolment **(A)** and onset **(B).** Kaplan-Meier curves for miR-181 (100 patients with subthreshold (blue) vs. 22 patients with suprathreshold (red) miR-181 levels) in replication cohort from enrolment **(C)** and onset **(D)**. Kaplan-Meier curves on the two cohorts merged (204 patients with subthreshold (blue) vs. 44 patients with suprathreshold (red) miR-181 levels) from **(E)** enrolment or **(F)** onset. Forest plot showing results of multivariate COX proportional hazard analysis of miR-181 corresponding to KM curves in panels A-F **(G).** *p<0.05, one-tailed Wald test. **p<0.01, one or two-tailed Wald test. ***p<0.001, two-tailed Wald test. Mean ± 95% CIs.

We next validated our results in an independent cohort of 122 patients, which was held-out until that point. Thus, we assessed discrimination between prognostic groups by miR-181, using the dichotomization miRNA threshold defined in the discovery cohort. Kaplan Meier curve analysis of plasma miR-181 levels in the replication cohort, also revealed clear survival curve separation between subgroups when survival was calculated from enrolment (logrank chi^2 =5.2, p=0.02, Figure 3C) or onset (logrank chi^2 =4.4, p=0.035, Figure 3D). Finally, we performed analysis on 248 patients in the combined cohort, from enrolment and from disease onset. Kaplan Meier analysis of plasma miR-181 levels in the combined cohort revealed clear survival curve separation between subgroups (enrolment, logrank chi^2 =18.5, p<0.0001, Figure 3E; onset, logrank chi^2 = 16.7, p<0.0001, Figure 3F). miR-181 levels were predictive of survival length, regardless of whether patients were treated with Riluzole or not (Figure S7).

Accordingly, COX regression analysis revealed significant hazard ratios from enrolment for high vs. low levels of miR-181 in the discovery cohort (HR 2.17, 95% CI: 1.25 - 3.75, p=0.006, Figure 3G), the replication cohort (HR 1.76, 95% CI: 0.97 - 3.18, one-tailed p=0.03), and the merged cohort (HR 2.09, 95% CI: 1.48 - 2.94, p<0.001). Likewise, hazard ratios, calculated from onset, were consistent for discovery (HR 2.83, 95% CI: 1.7 – 4.7, p<0.001), replication (HR 1.83, 95% CI: 1.1-3.0, one-tailed p=0.0087) and merged (HR 2.21, 95% CI: 1.56 – 3.12, p<0.001) cohorts.

In the discovery cohort, miR-181 displayed a 4-fold increase in patients with higher miR-181 levels compared to patients with low miR-181 levels (p<0.001, Figure S8A) while in the replication cohort, miR-181 levels increased by 8.5-fold in the high expression bin (p=0.009, Figure S8B). In addition, a modest but statistically significant correlation was found between plasma miR-181 levels and survival length from enrolment or onset (Figure S8C,D; Table S3). We further tested the D50 model-based descriptors, which is derivative of ALSFRS-R and addresses difficulties in characterizing aggression and the individual disease covered by traditional ALS clinical indices ^24^. Applying D50 to miR-181 stratification revealed association of high miR-181 levels with aggressive disease (time taken to reach half functionality < 32 months), whereas low miR-181 levels are associated with moderate disease (time to half functionality > 57 months, p<0.001, Fig. S9A; Table S3). Such a ∼ 25-month gap to losing half functionality might be clinically important. miR-181 levels also increased by 70% between mean value of patients suffering from aggressive (D50 < 45 months), relative to moderate (D50 > 45 months) disease (t-test: p=0.03, not shown).

miR-181 levels remain stable over time (Figure 1A, B), which is orthogonally supported by the lack of a difference in rD50, a measure of functional decline over the course of disease, between low and high miR-181 levels (p=0.07, Figure S9B), as well as the lack of correlation between miR-181 levels and rD50 (Figure S9C). Furthermore, miR-181 levels remained stable at early, progressive and late disease stages (0 ≤ rD50 < 0.25; 0.25 ≤ rD50 < 0.5; rD50≥0.5, respectively; ANOVA: p=0.15, Figure S9D). Therefore, miRNA measurements are unlikely to be biased by sampling at different disease stages. Finally, miR-181 levels were not correlated with progression rate, ALSFRS at enrolment or age at onset, and these clinical parameters were comparable between low and high miR-181 levels (Figure S10).

### miR-181 is broadly expressed in neurons

To elucidate the tissue source of miR-181 we revisited previously reported Nanostring data ^25^. miR-181a-5p is the ninth most abundant miRNA in laser capture micro-dissected human motor neurons of ALS patients and is also fairly abundant in the CNS in general ^26^. We further performed fluorescent *in situ* hybridization with a probe that hybridizes to miR-181a-5p in the mouse motor cortex and the lumbar spinal cord, two regions affected in ALS (Figure 4A, B). Punctate miR-181a-5p signal was found in motor cortex soma and neurites (Figure 4C) and in ventral horn neurons (Figure 4D). Thus, a conceivable source for miR-181 in ALS patients may be motor neurons in the cortex and spinal cord. The presence of miR-181 in neurites suggests that it could be an RNA marker of axonal damage, resembling the suggested axonal origin of protein biomarkers, such as NfL ^27^.

**Figure 4.**
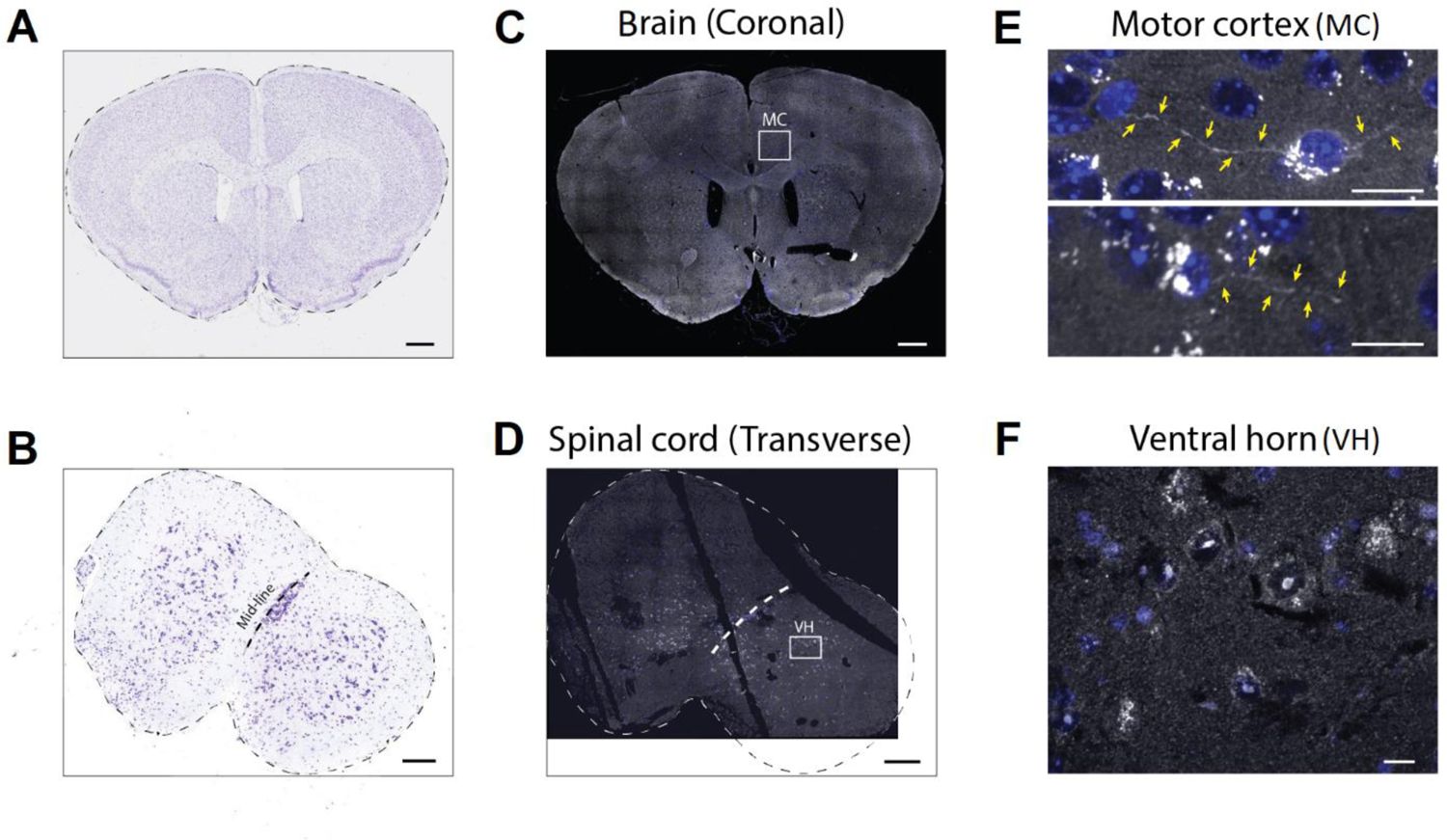
miR-181a-5p localizes to neuronal soma and neurites in mouse brain and spinal cord. Nissl images of mouse brain (**A**, 10X lens, scale bar 500μm) and lumbar spinal cord (**B**, magnification 10X, scale bar 200μm), and corresponding brain **(C)** and ventral horn **(D)** miR-181a-5p *in situ* hybridization micrographs. miR-181 is detected in critical motor neuron soma and neurites (**E**, motor cortex, 63X lens, scale bar 15μm), and in motor neurons of the spinal cord ventral horn (**F**, 63X lens, scale bar 20μm). MC: motor cortex; VH: ventral horn.

### miR-181 & NfL establish a cooperative miRNA-protein biomarker in prediction of ALS prognosis

We have previously shown that neurofilament light chain (NfL) can stratify ALS patients by their survival length ^28^. In the current cohort, we assayed NfL in all plasma samples with by single molecule array (Simoa) immunoassay. 243 of the 248 SIMOA samples were technically successful. A Cox proportional hazard analysis revealed that high plasma NfL predicts higher risk of death (from enrollment HR 2.09, 95% 1.49 – 2.94, p<0.001, C-index 0.59, Akaike’s information criteria (AIC) 2083, or from onset HR 2.26, 95% 1.73 – 2.96, p<0.001, C-index 0.62, AIC 2060 Figure 5A, B), as previously reported ^28–30^. The performance of miR-181 in predicting risk of death is comparable with that of NfL (from enrollment HR 2.03, 95% CI: 1.45 – 2.85, p<0.001, C-index 0.56, AIC 2096, or from onset HR 2.07, 95% CI: 1.6 – 2.7, p<0.001, C-index 0.56, AIC 2081, Figure 5A, B).

**Figure 5.**
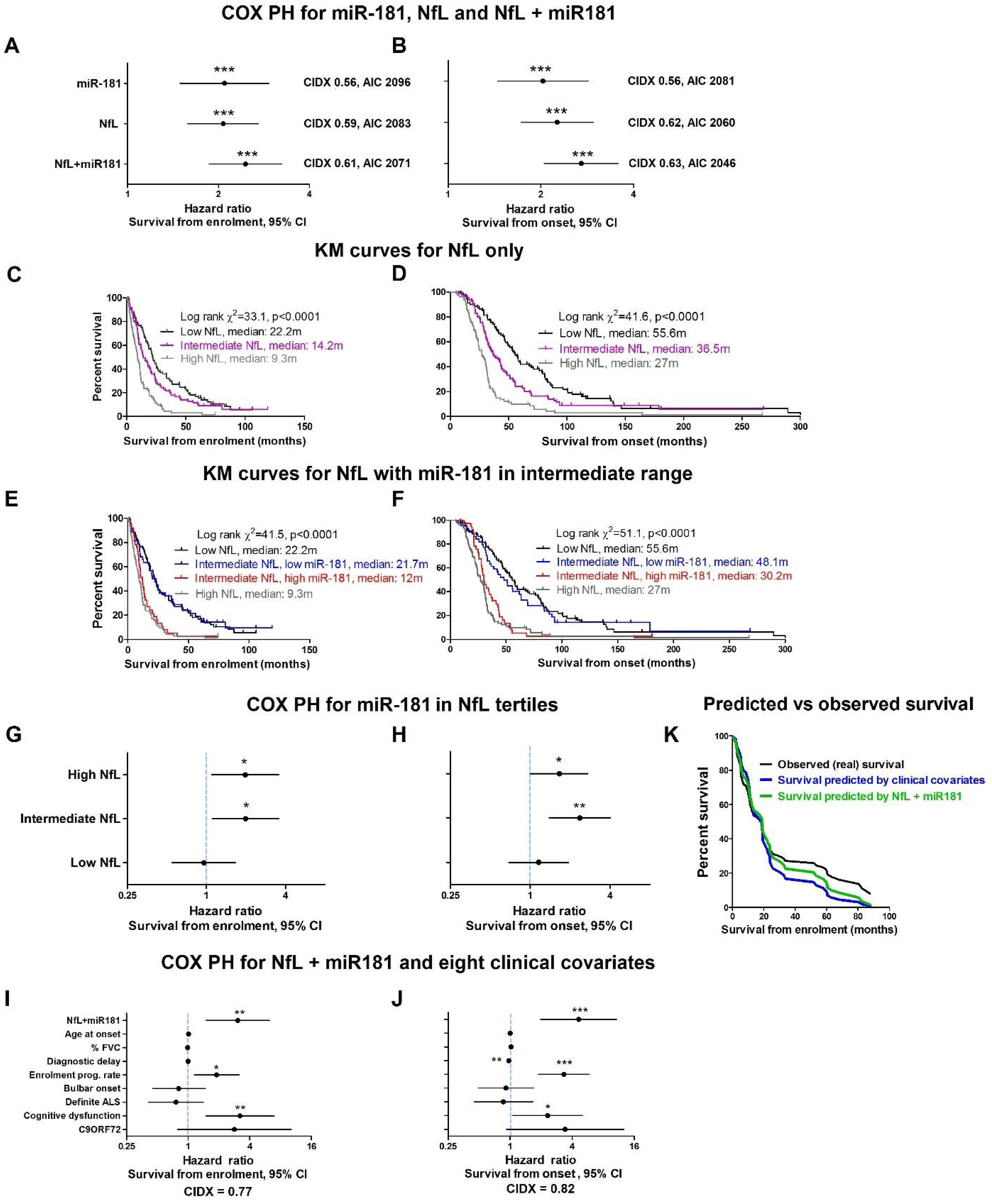
Superior accuracy for combination of miRNA and NfL biomarkers in prognosis analysis. Cox proportional hazard analysis for miR-181, NfL and a combinatorial predictor NfL+miR181 in 243 patients with both miR-181 and NfL measurements (threshold values: miR-181 71,000 UMIs, NfL 82.2 pg/ml, NfL+miR181 upper 118 vs lower 125), from enrolment **(A)** or onset **(B).** CIDX-concordance index. AIC - Akaike’s information criteria. Kaplan Meier curves calculated from enrolment **(C)** or onset **(D)** based on tertile stratification of plasma NfL levels. (NfL threshold values by Simoa assay: < 59 pg/ml (low); 59-109.8 pg/ml (intermediate); >109.8 pg/ml (high). Kaplan Meier curves calculated from enrolment **(E)** or onset **(F),** whereby the middle tertile of samples with intermediate NfL levels is further subdivided by miR-181 levels. Forest plots of mortality hazard ratio, calculated by survival length from enrolment **(G)** or onset **(H)** for high vs low miR-181 levels in the three NfL tertiles. Forest plots of mortality hazard ratio, calculated by survival length from enrolment **(I)** or onset **(J)** for combined miRNA-protein predictor, NfL+miR181 and 8 clinical covariates ^23^ on a subset of 75 patients. Mean ± 95% CI, Wald test *p<0.05, **p<0.01, ***p<0.001. **(K)** Observed survival curve (black) vs. prediction based on NfL + miR181 (green) or by 8 clinical covariates (blue), in a subset of 75 patients.

We then tested a combined predictor based on both NfL and miR-181, creating a binary protein-miRNA feature “NfL+miR181”. An interaction variable based on both NfL and miR-181, yielded higher risk of death than each one of single markers on its own (from enrolment, HR 2.46, 95% CI: 1.87 – 3.24, p<0.001, C-index 0.61, AIC 2071, or from onset: HR 2.7, 95% CI: 2.05 – 3.56, p<0.001, C-index 0.63, AIC 2046, Figure 5A, B). Therefore, miR-181 and NfL display comparable capacities, as single estimators of death risk, in patients with ALS. However, together the miRNA-protein pair displays a cooperative predictive value. We then stratified samples into tertiles, according to NfL levels, which exhibits different survival length ^28^, (Figure 5C, D). Interestingly, in the range of intermediate NfL levels, the additional stratification by miR-181 separated this sub-cohort in two, as revealed by KM analysis (from enrollment logrank chi^2 = 41.5, p<0.0001, from onset, logrank chi^2 = 51.1, p<0.0001, Figure 5E, F). Cox regression analysis on miR-181 levels in the low, intermediate, and high NfL tertiles revealed a higher risk of dying with higher miR-181 plasma levels in the middle and high tertiles (from enrolment: HR 2.0, 95% CI, 1.1 - 3.6, p=0.03, Figure 5G), but not in the low tertile (HR 0.96, 95% CI, 0.55 - 1.66, p=0.9). Similarly, when calculated from disease onset, higher miR-181 levels predicted a higher risk of dying for patients within the range of intermediate NfL tertile (HR 2.37, 95% CI, 1.4-4.02, p=0.001, Figure 5H) and a modest added risk in the high NfL tertile (HR 1.66, 95% CI, 1.0-2.7, p=0.04). Therefore, miR-181 may be valuable in particular at the range of intermediate NfL values, where it can accurately call a 18 months difference in median prognosis that cannot be identified by measurements of NfL alone.

We tested the potential correlation of miR-181 with other molecular biomarkers that are under investigation, neurofilaments, TNF, creatinine and creatine kinase in the same cohort. Notably, miR-181 levels did not correlate with the levels of other plasma biomarkers, (Figure S11; Table S3), suggesting it works via an alternative mechanism.

Finally, we were interested in the relationship of bimolecular blood predictors and established clinical features of the disease. Thus, we performed multivariate Cox analysis using the combined predictor NfL+miR181 along with eight other clinical features that were previously shown to be informative (age of onset, forced vital capacity, diagnostic delay, enrolment progression rate, site of onset, El Escorial’s definite ALS, cognitive dysfunction and C9orf72 genetics)^23^. High NfL+miR181 predicted a risk of dying that was 3-4.6 times higher (from enrolment HR 3.06, 95% CI: 1.5 - 6.24, p = 0.002; from onset HR 4.63, 95% CI: 1.98 - 10.82, p < 0.001, Figure 5I, J). We have also performed a digital PCR study to quantify miR-181 RNA molecule concentration in human plasma. Unique molecular identifiers (UMIs) in sequencing correlated to absolute miRNA copies by digital PCR with Pearson R^2^ 0.97 (327, 389, 523, 688 UMIs corresponding to 4020, 5760, 6960, 9540 miR-181 RNA molecules /microliter of human plasma). This analysis further suggests that the threshold of miR-181, when utilized with NfL as biomarker pair, is at approximately 5340 RNA molecules /microliter of human plasma).

Finally, the predicted survival curve, with the miRNA-protein predictor NfL+miR181, is closer to the observed (real) survival curve, than survival curve approximated by the multivariate cox model with eight established clinical features. Together, miR-181 stands on its own as a powerful prognostic marker for ALS. Furthermore, utilization of miR-181 in concert with an established protein biomarker, NfL, is more accurate than either alone.

## Discussion

In this study, we report the results of one of the most elaborated small RNA-seq studies, undertaken to date in neurodegeneration research. We show that in ALS, miRNAs appear to be mostly unchanged longitudinally during disease (Figure 1), whereas increase in miR-423/484/92a/b levels during disease course could contribute to monitoring of disease progression (Figure S2,3).

Importantly, high miR-181 levels predicted shortened survival in two ALS cohorts. miR-181 is encoded from a human gene that has seen local duplication to a bi-cistronic miR-181a and miR-181b within the same transcriptional unit and additional genomic duplications that results in three homologs across the human genome. miR-181a and miR-181b are functionally identical, silence the same target set, and are co-expressed from the same gene. Although equivalent, the simultaneous consideration of both RNAs provides superior sensitivity as a predictor of ALS prognosis and progression (Figure 2). miR-181 ability to predict prognosis of patients with ALS was validated in a replication cohort (Figure 3). The fact that miR-181 levels stay stable during the course of disease, suggests a constant process underlying their generation and clearance rates. miR-181 species are expressed in the brain and spinal cord, including in cortical and spinal motor axons and soma (Figure 4) and their transport and biogenesis is regulated in neuronal axons ^31^. Therefore, it may be that the utility of miR-181 as a prognostic biomarker in ALS is linked to being spilled off dying axons, somewhat reminiscent of NfL, which is a neuronal cytoskeletal protein. Accordingly, we demonstrate that miR-181 and NfL serve separately as predictors of ALS prognosis, with comparable predictive capacity (Figure 5). Furthermore, miR-181 measurement can enhance the prognostic value of NfL and a joint miRNA-protein measure may compute prognosis more precisely than any of the circulating biomolecules on their own. Specifically, we show that miR-181 levels were of predictive value particularly when NfL values are intermediate, and the combination of miR-181 and NfL is able to discriminate fast and slow progressors in this group. Stratification based on progression rate is important in clinical trials to balance treatment and placebo groups. Indeed, certain trials have focused on ALS fast progressors in order to obtain outcomes that are more reliable. Our proposed combination of miR-181 and NfL as a prognostic biomarker would enhance our ability to predict ALS progression and thereby facilitate recruitment to clinical trials. Therefore, the development of a miR-181-based biomarker opens a new horizon for a combinatorial protein-RNA biomarker system for ALS prognosis and encourages testing the value of orthogonal multi-omic platforms for additional biomarker endpoints.

We also found miR-181 expressed in the brain and the spinal cord, and suggest that plasma miR-181 originates in part from the central nervous system, reminiscent of NfL. While miR-181 was reported in the cerebrospinal fluid of ALS patients, and might contribute to ALS diagnosis ^19^, it is also abundant in hematopoietic tissues ^32^, which also contribute to its presence in the plasma. That miR-181 levels do not correlate with clinical features or several other circulating biomolecules may perhaps reflect different facets of the medical condition or disparate underlying mechanisms.

Several additional factors contribute to the potential clinical impact miR-181 quantification. First, blood immune-complexes and antibodies against NfL rise with disease progression, which will likely limit the dynamic range of NfL-based predictors under some circumstances ^33^. In such cases, the value of miR-181 may be even more pronounced. In addition, real-world patient profiles indicate that most ALS patients are in the middle tertile neurofilament level, ^30^ the enhanced sensitivity provided by miR-181 at this region of the spectrum will be particularly useful. Taken together, miR-181 emerges as a prognostic ALS biomarker that can be developed in combination with NfL to improve the accuracy of patient stratification in clinical trials.

## Online Methods

### Standard protocol approvals, registrations, and patient consents

The study included a cohort with 252 patients with ALS from the ALS biomarker study. Patients were diagnosed with ALS according to standard criteria by experienced ALS neurologists ^34^. All participants provided written consent (or gave verbal permission for a carer to sign on their behalf) to be enrolled in the ALS biomarkers study if they met inclusion criteria until the desired sample size was reached (consecutive series). Ethical approval was obtained from East London and the City Research Ethics Committee 1 (09/H0703/27).

### Study design

We determined the sample size by doubling this number calculated by the following power analysis: 120 ALS patients are needed to obtain a hazard ratio of 3 with a power of 99% and a p-value of 0.01. A full cohort of 252 patients was randomly split into a discovery and replication cohorts with comparable clinical characteristics, each with 126 patients. Phenotypic data on de-identified patients was separated and blinded during steps of the molecular analysis. Disease severity was assessed with the revised ALS Functional Rating Scale (ALSFRS-R)^35^, and progression rate at enrolment (i.e. first blood draw) was calculated as follows: (48 - enrolment ALSFRS-R)/time (in months) from symptom onset to enrolment. Progression was also modeled using the D50 model which fits a sigmoid decay across all available ALSFRS-R scores ^36, 37^. Use of Riluzole (or not) at the time of sampling was recorded. Blood was collected by venipuncture in EDTA tubes, and plasma was recovered from the whole blood sample by centrifugation and stored at −80°C until performing downstream assays (RNA-seq and SIMOA for NfL).

### Longitudinal cohort analysis

Serial plasma samples and clinical information were obtained, on average, every 2 to 4 months from 48 patients with ALS. No selection criteria were applied to individuals with ALS sampled longitudinally, other than their willingness to donate further samples. Longitudinal analysis of miRNAs was first performed on samples from 22 patients, in an unbiased manner by next generation RNA sequencing. Results were tested on a replication longitudinal cohort of 26 patients by an orthogonal method of quantitative real time PCR. Symptom onset was defined as first patient-reported weakness.

### Small RNA next generation sequencing

Total RNA was extracted from plasma using the miRNeasy micro kit (Qiagen, Hilden, Germany) and quantified with Qubit fluorometer using RNA broad range (BR) assay kit (Thermo Fisher Scientific, Waltham, MA). For small RNA next generation sequencing (RNA-seq), libraries were prepared from 7.5 ng of total RNA using the QIAseq miRNA Library Kit and QIAseq miRNA NGS 48 Index IL (Qiagen), by an experimenter who was blinded to the identity of samples. Samples were randomly allocated to library preparation and sequencing in batches. The longitudinal ALS study samples were sequenced in one batch to avoid batch-induced biases in interpretation of longitudinal changes (analyzed in Figure 1). Precise linear quantification of miRNA is achieved by using unique molecular identifiers (UMIs), of random 12-nucleotide after 3’ and 5’ adapter ligation, within the reverse transcription primers ^20^. cDNA libraries were amplified by PCR for 22 cycles, with a 3’ primer that includes a 6-nucleotide unique index, followed by on-bead size selection and cleaning. Library concentration was determined with Qubit fluorometer (dsDNA high sensitivity assay kit; Thermo Fisher Scientific, Waltham, MA) and library size with Tapestation D1000 (Agilent). Libraries with different indices were multiplexed and sequenced on NextSeq 500/550 v2 flow cell or Novaseq SP100 (Illumina), with 75bp single read and 6bp index read. Fastq files were de-multiplexed using the user-friendly transcriptome analysis pipeline (UTAP) ^38^. Human miRNAs, as defined by miRBase ^39^, were mapped using Geneglobe (Qiagen). Sequencing data normalized with DESeq2 package ^40^ under the assumption that miRNA counts followed negative binomial distribution and data were corrected for the library preparation batch in order to reduce its potential bias.

### Selecting candidate miRNA and miRNA pairs for prognostic analysis

The pipeline is succinctly described in Figure S5. 2008 miRNAs were aligned to the genome in the longitudinal study and out of them, 187 miRNAs that exhibited >50 UMI counts in 60% of the samples and non-zero counts in all samples, were included in further analysis. 125 out of these 187 miRNAs were longitudinally stable with low inter-individual variability (blue features in Figure 1A). In the discovery cohort, 106 out of these 125 miRNAs passed a filtering criterion of average UMI counts >100 across all samples and non-zero counts, and were analyzed for prognosis differences between low and high level in the discovery cohort. Then, the miRNAs predictors were transformed from a continuous expression level to binary predictors (high/low), when the optimal dichotomization cut-off values were determined by iterative logrank analysis on all possible sample distributions for the remaining 106 miRNAs ^21^. 19 miRNAs were further excluded after additional QC if different miRNAs of the same family provided conflicting prognosis predictions, e.g. miR-27a predicted beneficial prognosis and miR-27b a detrimental prognosis). A logrank test, to compare survival distributions was performed for the remaining 87 miRNAs and null hypothesis significance testing (p values) for prognosis differences as demonstrated in Figure 2. Nine out of 87 miRNAs displayed logrank p ≤0.01. All 36 combinatorial pairs of these 9 miRNAs (9*8/2=36), were also subjected to logrank test to test cooperative prediction, via multiplication of the levels of two single miRNAs and then transforming them to a binary (high/low) predictors, as done for single miRNA predictors. 20 (out of 36) miRNA pairs displayed logrank p ≤0.01. A total of 29 candidate prognostic biomarkers (9 miRNAs and 20 miRNA pairs) were then examined under bootstrap feature selection.

### Model selection: AIC-based backward feature selection by bootstrap resampling

Feature selection by bootstrap resampling was performed on a full cox model of 29 features (9 single candidate miRNAs and 20 miRNA pairs) using stepwise backward elimination based on Akaike information criterion (AIC) ^22^. The cox regression coefficients and standard errors are estimated in the full model (null model), including all variables under consideration, and at each step a single feature is eliminated until no significant improvement in AIC is obtained. The procedure is repeated for 100 bootstrap samples, that were randomly drawn from the original cohort (n=126). Within each bootstrap sample, a cox model is developed and a list of selected features that optimize AIC is obtained. Candidate biomarkers are then ranked according to the proportion of bootstrap samples in which they were selected as best predictors, and the proportion of bootstrap samples where their cox coefficient was significant (at significance level 0.05). We considered the following criteria for selecting the final prognostic biomarkers from the bootstrap resampling procedure: features that were selected >70% of bootstrap samples and were statistical significance in >85% of the bootstrap samples in which they were selected.

Only a single predictor fulfilled these criteria, miR-181. A univariate cox model of miR-181, stratified by ALSFRS slop and age of onset is then assessed on discovery and replication cohort. A numerical threshold of 71,000 UMIs, which was found as optimal by Evaluate Cutpoints ^21^ in the discovery cohort (N=126), separated between sub-threshold patients (N=104) and supra-threshold patients (N=22). Same threshold was used in an independent replication cohort, whereby 4 out of 126 samples, with borderline miR-181 levels, were excluded. Joint analysis with the same threshold was further conducted on the combined cohort of 248 patients.

### Polymerase chain reaction assays

Quantitative real time PCR (qPCR) of miR-423/484/92a/92b, performed with Taqman advanced miRNA assay probes (Thermo Fisher) with the following probes: hsa-miR-423-5p (Assay ID: 478090_mir); hsa-miR-484 (Assay ID: 478308); hsa-miR-92a-3p (Assay ID: 477827); hsa-miR-92b-3p (Assay ID: 477823). hsa-miR-140-3p (Assay ID: 477908) and hsa-miR-185-5p (Assay ID: 477939) were selected as normalizers, based on stable plasma levels in the longitudinal cohort, described in Figure 1: (1) basemean expression between 500-3,000; (2) coefficient of variation ≤ 0.35.cDNA Synthesis Kit (Applied Biosystems) was used for cDNA reverse transcription (10 ng input) and run on a StepOnePlus (Applied Biosystems). Data compared between samples at enrolment (t_1_) and corresponding follow-up sample (t_2_) by one-tailed paired t-test. Digital droplet PCR (ddPCR) was performed using the hsa-miR-181a and hsa-miR-181b probes (Taqman assay ID: 000480, 001098, respectively, Thermo Fisher Scientific). Mix (5 μL of cDNA 11 μL ddPCR supermix (Bio-Rad), 1 μL ×20 TaqMan Assay, 5 μL H_2_O) was gently vortexed, droplets were generated in QX100/QX200 with DG8 cartridges (Bio-Rad) and put into 96-well in C1000 thermocycler (Bio-Rad) for a protocol: 95°C, 10 minutes (1 cycle), 60°C annealing/extension step, 1 minute followed by 94°C melting step, 30 seconds (39 cycles), and a final stage of 98°C, 10 minutes followed by holding at 12°C. Plates read on the QX200 droplet and analyzed by QuantaSoft software (Bio-rad) after setting a FAM threshold based on the ‘no template’ negative control fluorescence histogram.

### Neurofilament light chain (NfL) assay

The quantitative determination of NfL in human plasma was undertaken by Single Molecule Array technology using a digital immunoassay Simoa HD-1 Analyzer (Quanterix, Lexington, MA). Standards, primary and secondary antibodies, detection range including lower and upper limits of detection were specified by manufacturer (Simoa Nf-L Advantage Kit-102258, Quanterix). An equal volume was loaded for all samples in study. NfL threshold concentrations were defined by cohort tertiles: *NfL <59 pg/ml* for the lower tertile *(81 patients), 59-109.8 pg/ml* for the middle tertile *(81 patients), or >109.8 pg/ml* for the higher tertile *(81 patients)*.

### RNA *in situ* hybridization

Mouse studies, performed in accordance with institutional guidelines and IACUC. Adult mice were deeply anesthetized (10% ketamine, 2% xylazine in PBS, 0.01 ml / gram body weight) and intracardially perfused with 10ml of PBS, followed by 40ml of 4% Paraformaldehyde. Brains and spinal cords were dissected, fixed in fresh 4% PFA at room temperature for 24 hrs., dehydrated in graded ethanol series, cleared with ethanol / histoclear (1:1 vol. / vol.) and then in histoclear (National Diagnostics) and embedded in paraffin. 4 µm microtome sections were mounted onto Superfrost plus slides and deparaffinized. miRNA *in-situ* hybridization performed with hsa-miR-181a-5p probe (VM1-10255-VCP, ViewRNA Tissue Assay, Thermo-Fisher Scientific), counterstained with DAPI and mounted with ProLong Gold (Molecular Probes, P36934). Adjacent sections were taken for cresyl violet (Nissl) staining. Micrograph acquisition performed on Dragonfly Spinning disc confocal system (Andor Technology PLC) with Leica Dmi8 Inverted microscope (Leica GMBH) with 10X (air) and 63X (glycerol) objectives equipped with sCMOS Zyla (Andor) 2048X2048 Camera. DAPI (Excitation 405nm, emission 450/50nm, 100ms); ViewRNA probe (excitation 561nm emission 620/60nm, 200ms). Background and Shading Correction was performed using BaSIC ^41^.

### Analysis of the combination of NfL and miR-181 as prognostic factors

Logical operators developed to define a new combined miRNA-protein predictor for ALS prognosis:

If (*NfL < 59 pg/ml) or (NfL 59-109.8 pg/ml AND miR-181 < 39,300* UMIs*) = NfL+miR181 = 0* If (*NfL >109.8 pg/ml) or (NfL 59-109.8 pg/ml AND miR-181 > 39,300* UMIs*) = NfL+miR181 = 1*

### Analysis of clinical features as prognostic factors

A subset of 75 patients out of the all 252 participates of the study, held the complete clinical information sufficient to perform multivariate COX analysis with *NfL+miR181*eight clinical covariates (diagnostic delay, forced vital capacity, C9ORF72 genetics, progression rate at enrolment, cognitive dysfunction, age at onset, bulbar onset and definite ALS by El-Escorial criteria) that were described in ^23^.

### Statistical analysis

In longitudinal cohort, p values were calculated by Wald test ^40, 42^ and adjusted for multiple testing according to Benjamini and Hochberg ^43^. Logrank Mantel-Cox test was used for Kaplan-Meier survival estimators and a fixed date was used to censor data for survival analysis. Optimal dichotomization cut-off values of miRNA levels determined using Evaluate Cutpoints ^21^. Multivariate or univariate Cox proportional hazard analyses were used to calculate mortality hazard ratios, with molecular and phenotypic features as covariates. For longitudinal miRNA expression by qPCR, one-tailed paired t-test was used. Outliers were detected by Grubbs test ^44^ and excluded from analysis. Tests were run in R Project for Statistical Computing environment ^45^ and graphs were generated with Prism 5 (GraphPad Software, San Diego, California, USA).

## Acknowledgements

We thank Vittoria Lombardi (UCL) for technical assistance, and Iddo Ben-Dov from Hadassah Hebrew University Medical Center, Jerusalem, Israel, for advice on statistics. We acknowledge patients for their contribution and all the ALS biomarkers study co-workers and their contribution to the biobanking project, which has made this study possible (REC 09/H0703/27). We also thank the North Thames Local Research Network (LCRN) for its support and life science editors for editorial assistance. EH is the Mondry Family Professorial Chair and Head of the Nella and Leon Benoziyo Center for Neurological Diseases. Imaging performed at de Picciotto Cancer Cell Observatory, in memory of Wolfgang and Ruth Lesser.

## Funding

This research was supported by the following grants: Motor Neuron Disease Association (MNDA #839-791). Research at Hornstein lab is supported by CReATe consortium and ALSA (program: ‘Prognostic value of miRNAs in biofluids from ALS patients’), RADALA Foundation; AFM Telethon (20576); Weizmann - Brazil Center for Research on Neurodegeneration at Weizmann Institute of Science; the Minerva Foundation with funding from the Federal German Ministry for Education and Research, ISF Legacy Heritage Fund 828/17; Israel Science Foundation 135/16; 392/21; 393/21; 3497/21. Target ALS 118945; Thierry Latran Foundation for ALS Research. the European Research Council under the European Union’s Seventh Framework Programme (FP7/2007–2013)/ERC grant agreement number 617351; ERA-Net for Research Programmes on Rare Diseases (eRARE FP7) via Israel Ministry of Health; Dr. Sydney Brenner and friends, Edward and Janie Moravitz, A. Alfred Taubman through IsrALS, Yeda-Sela, Yeda-CEO, Israel Ministry of Trade and Industry; Y. Leon Benoziyo Institute for Molecular Medicine, Kekst Family Institute for Medical Genetics; David and Fela Shapell Family Center for Genetic Disorders Research; Crown Human Genome Center; Nathan, Shirley, Philip and Charlene Vener New Scientist Fund; Julius and Ray Charlestein Foundation; Fraida Foundation; Wolfson Family Charitable Trust; Adelis Foundation; Merck (United Kingdom); M. Halphen; and Estates of F. Sherr, L. Asseof, L. Fulop. From:benatPF is supported by an MRC/MND LEW Fellowship and by the NIHR UCLH BRC. JG was supported in the JPND framework ONWebDUALS and LG is the Graeme Watts Senior Research Fellow supported by the Brain Research Trust. NSY was supported by the Israeli Council for Higher Education (CHE) via the Weizmann Data Science Research Center. IM was supported by Teva Pharmaceutical Industries Ltd. as part of the Israeli National Network of Excellence in Neuroscience (NNE, fellowship 117941).

## Competing interests

The authors state that they have no competing interests.

## Data availability

Source data for figures are provided in supplementary tables. Fastq.gz files with raw sequencing data, text files with raw read counts, excel files with processed read counts and R codes are available as <u>GSE 168714</u> in gene expression omnibus (GEO).

---

**Figure S1.**
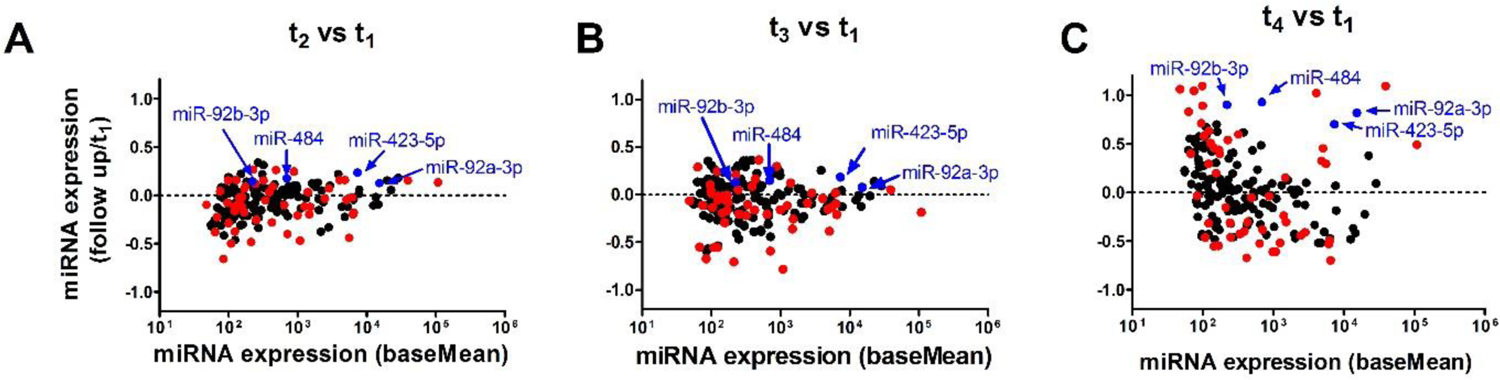
(A-C) MA plots of differential miRNA expression upon repeated sampling relative to the first phlebotomy. Red features denote miRNAs with statistically significant change in levels.

**Figure S2.**
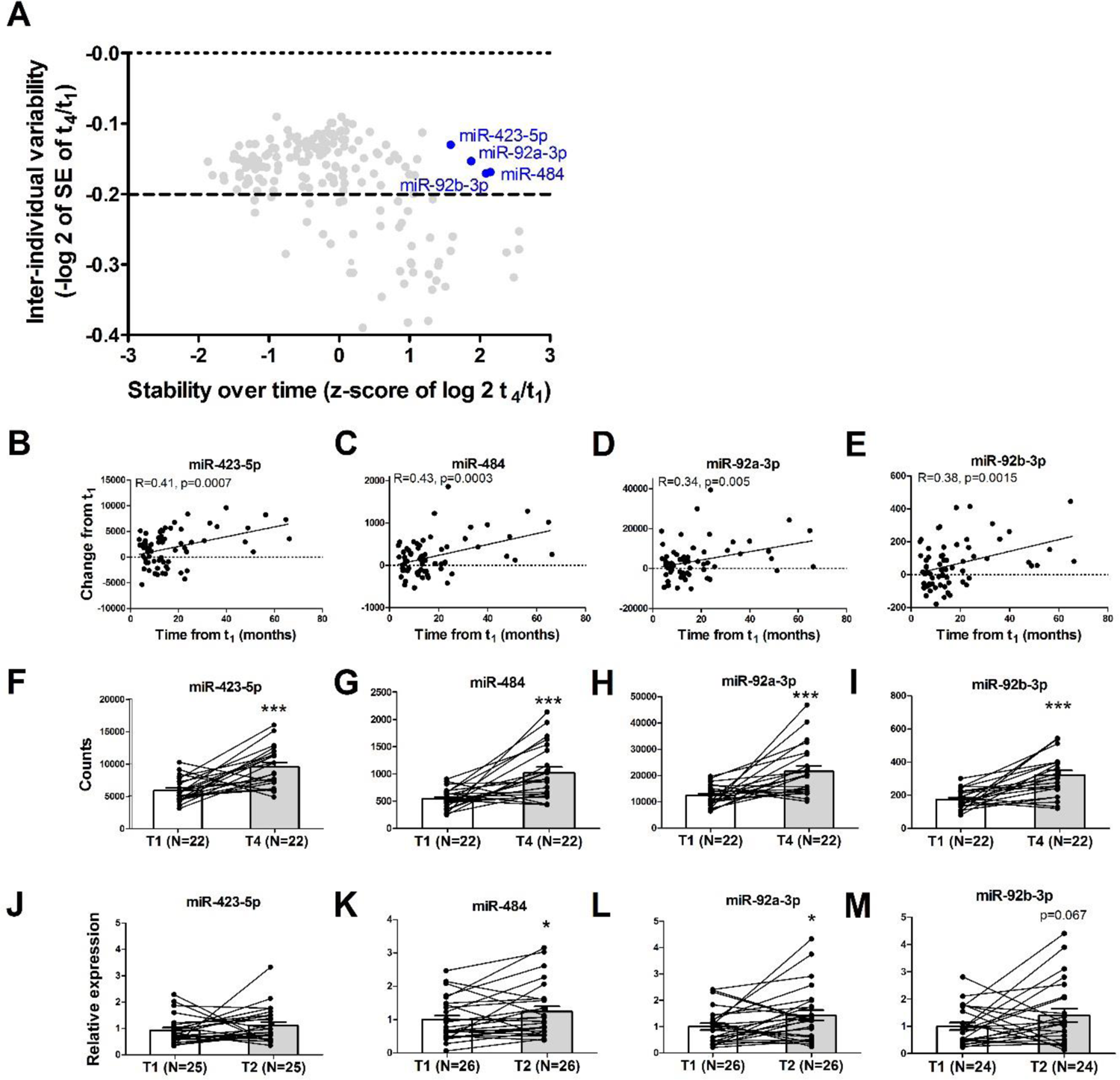
Analysis of miRNAs that increase during ALS course. Plasma levels of four miRNAs of the 129 miRNAs analyzed in main Figure 1A displayed low inter-individual variability, but increased with the disease course, suggesting that, although they are not suited for prognostic use, they could potentially monitor disease progression. (miR-423/484/92a/b, t_4_/t_1_ > 1.5 SD, X-axis) **(A)**. Temporal changes in the levels of **(B)** miR-423-5p, **(C)** miR-484, **(D)** miR-92a-3p, or **(E)** miR-92b-3p and revealed correlation with time passing from enrolment (in months). Spaghetti plots of individual patient trajectories (t_1_-t_4_) denoted for **(F)** miR-423-5p, **(G)** miR-484, **(H)** miR-92a-3p, or **(I)** miR-92b-3p. Time intervals: t_1_-t_2_ 6.3±0.3 m.; t_1_-t_3_ 13.0±0.3 m.; t_1_-t_4_ 32.7±3 m. Disease duration: t_1_ 28.8±3 m.; t_4_ 61.5±3 m. Validation of changes to miRNA levels in an independent replication cohort (N=26 individuals, Table 2). Spaghetti plots of individual patient trajectories (t_1_-t_2_ 13.7 ± 1.6 months) in a replication cohort, for **(J)** miR-423-5p, p=0.17 **(K)** miR-484, p=0.02; **(L)** miR-92a-3p, p=0.02; or **(M)** miR-92b-3p, p=0.067. One-tailed t-test. Together, miR-484 and miR-92a/b may be considered as candidate molecular biomarkers of functional decline over the course of disease. Mean ± SEM. *p<0.05, ***p<0.001, one-tailed paired t-test or Wald test. Analysis of a single miR-423-5p sample and two miR-92b-3p samples in the replication cohort deviated from the mean according to Grubbs test and these were excluded as outliers.

**Figure S3.**
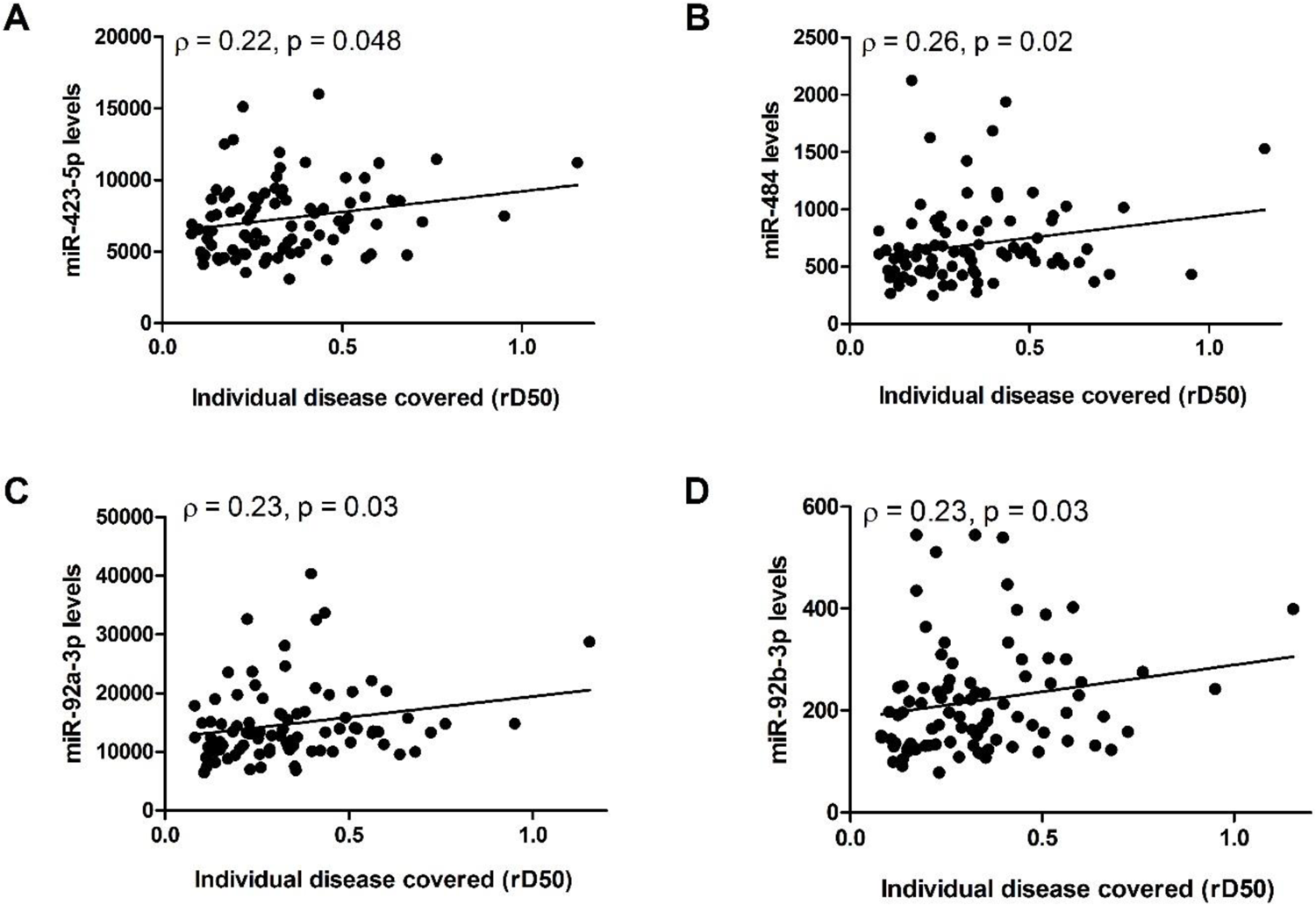
Longitudinal change of miRNA-423-5p, miR-484, miR-92a and miR-92b with disease progression. Correlation between the relative disease covered (rD50) in longitudinal plasma collections (X-axis) and levels of **(A)** miR-423-5p, **(B)** miR-484, **(C)** miR-92a-3p, and **(D)** miR-92b-3p. The relative D50 (rD50) is a derivative of ALS Functional Rating Scale-Revised (ALSFRS-R) decay that reveals the disease covered by individual patients independent of the rate of progression ^24, 37^. For example, an rD50 of 0.0 signifies ALS onset, and 0.5 signifies the time-point where functionality is reduced by half. Longitudinal miR-484/92a/b levels in blood correlated with rD50 at the time of sampling (Figure S3A-D).

**Figure S4.**
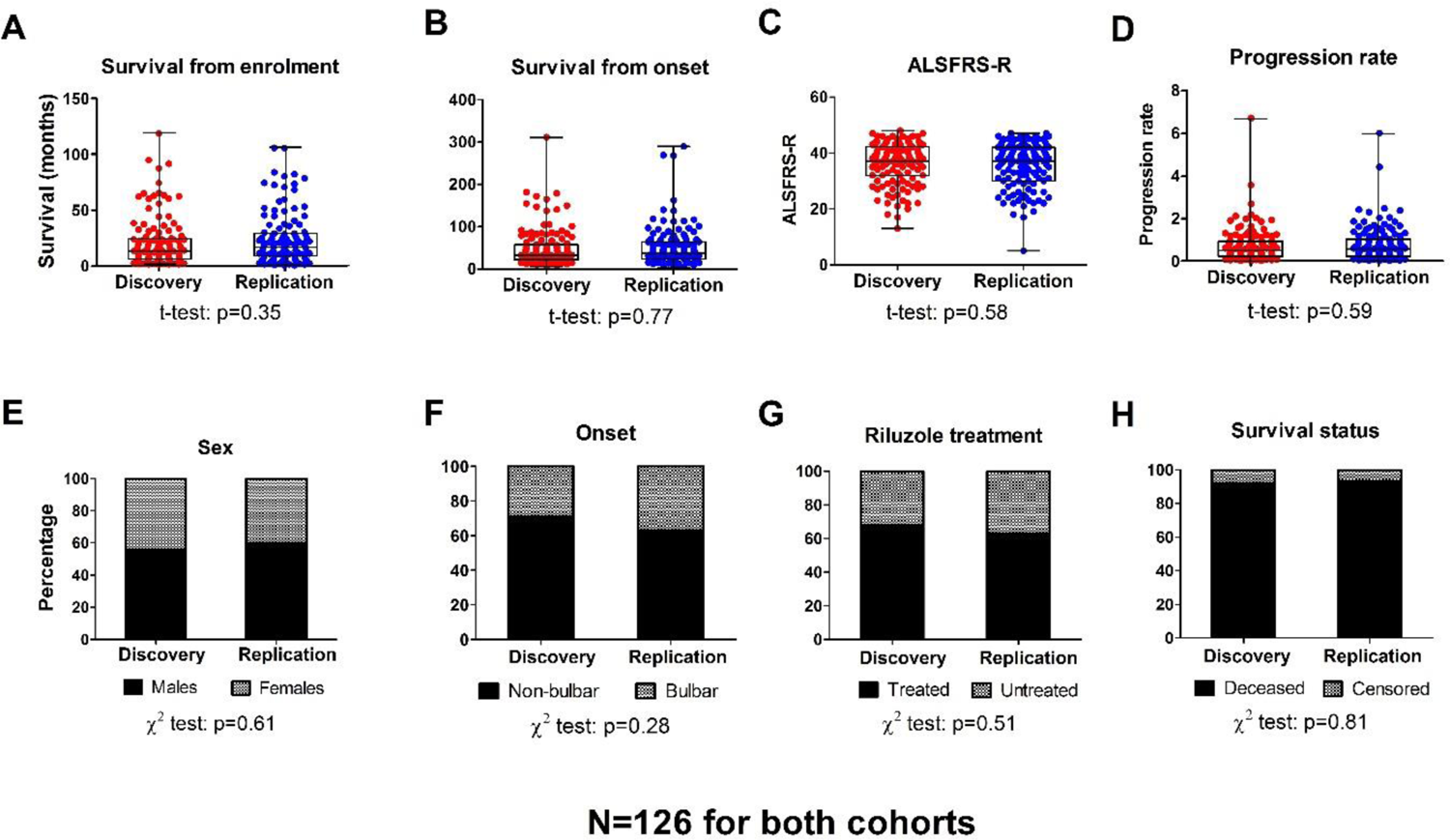
Clinical features are comparable between cohort I and cohort II. (A) Survival from enrolment **(B)** survival from symptom onset **(C)** ALSFRS-R score at enrolment **(D)** progression rate at enrolment **(E)** sex distribution **(F)** onset site distribution **(G)** Riluzole treatment status **(H)** number of censored patients.

**Figure S5.**
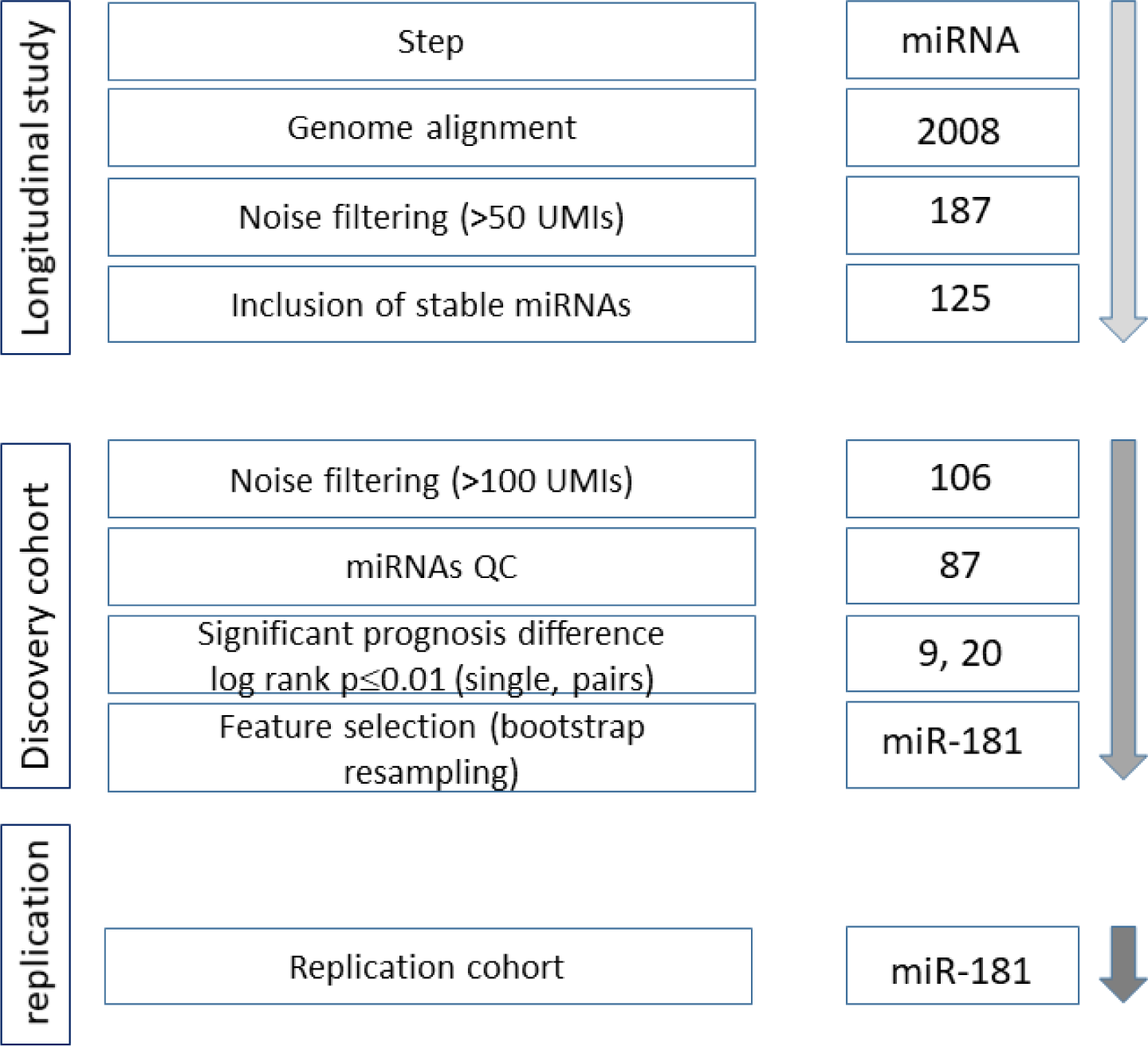
Pipeline for selecting miRNAs as candidate prognostic markers. 2008 miRNAs were aligned to the genome in the longitudinal study and out of them, 187 miRNAs which exhibited >50 UMI counts in 60% of the samples were included in further analysis. 125 out of the 187 miRNAs were longitudinally stable with low interindividual variability (green features in Figure 1A). In the discovery cohort, 106 out of these 125 miRNAs passed a filtering criterion of average UMI counts >100 across all samples, and were analyzed for prognosis differences between low and high level in the discovery cohort. 19 miRNAs were further excluded after additional QC based on logrank analysis (opposite directions of prognosis differences between members of the same miRNA family, e.g. miR-27a and miR-27b), and the remaining 87 miRNAs were assessed for logrank and p values for prognosis differences as demonstrated in Figure 2. 9 out of these 87 miRNAs displayed logrank p≤0.01, and all of their possible pairs (9*8/2=36), derived from multiplication of the levels of two single miRNAs, were further assessed for prognosis differences. Nine single miRNAs and 20 miRNA pairs which displayed logrank p≤0.01 were further subjected to feature selection by bootstrap resampling, whereby features had to be selected >70% of the bootstrap samples and display statistical significance in >85% of the samples in which it selected, in order to be tested as a prognostic marker on the full discovery cohort. miR-181 was the only feature fulfilling those criteria, hence it was tested in the discovery cohort, and exhibited significant survival differences and hazard ratios, both in the discovery cohort and when validated on a replication cohort that was set aside until that point.

**Figure S6.**
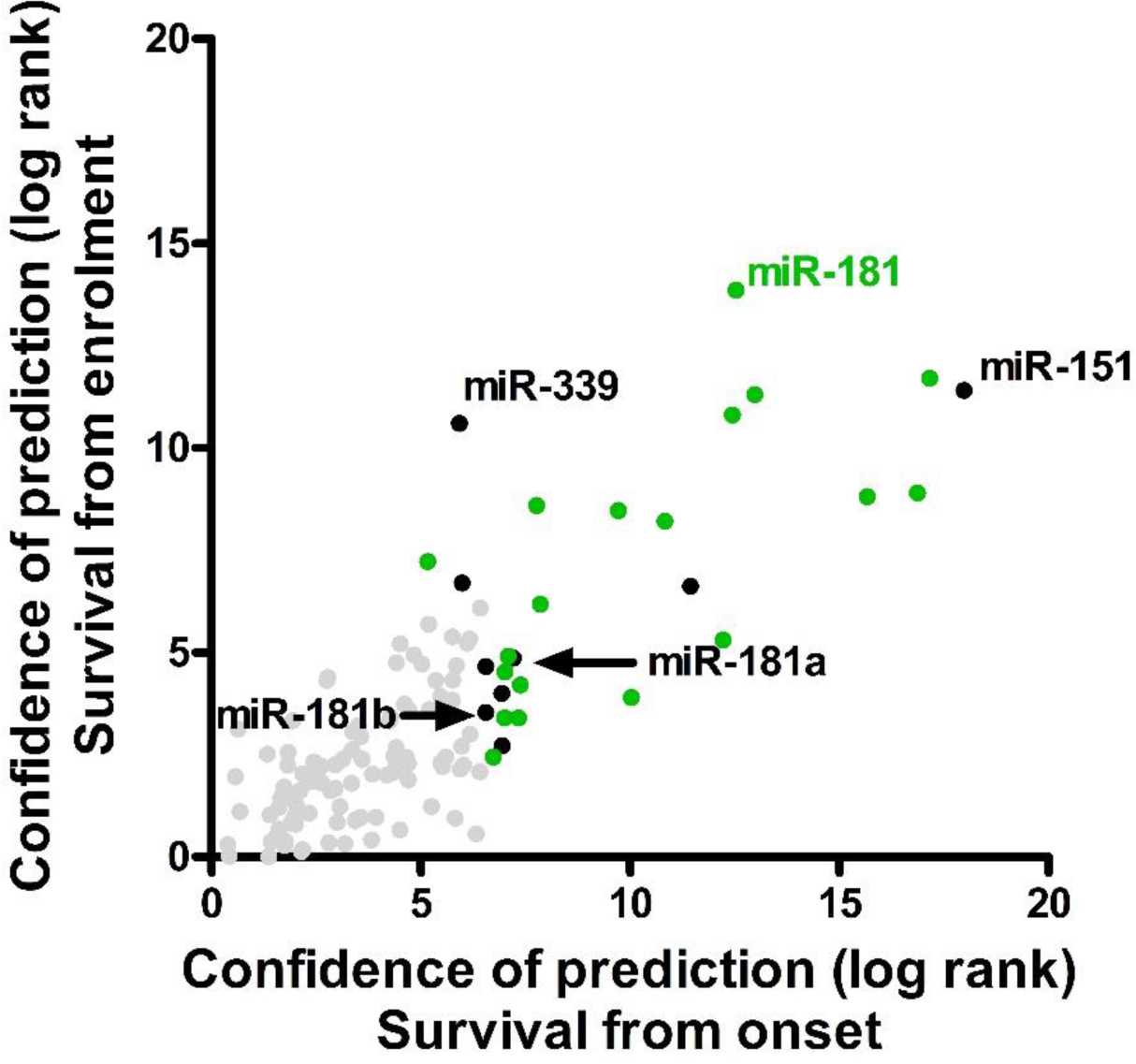
Scatter plot, assessing agreement between separation of survival by 123 miRNA features, by logrank test from study enrollment (Chi^2, y-axis) or first symptoms (onset, Chi^2, x-axis). The optimal threshold was calculated per miRNA in a discovery cohort of 126 patients by^21^. Single miRNA (black) or miRNA pairs (green), displaying a p-value ≤0.01 (log 10 transformed values ≥ 2), and grey: insignificant.

**Figure S7.**
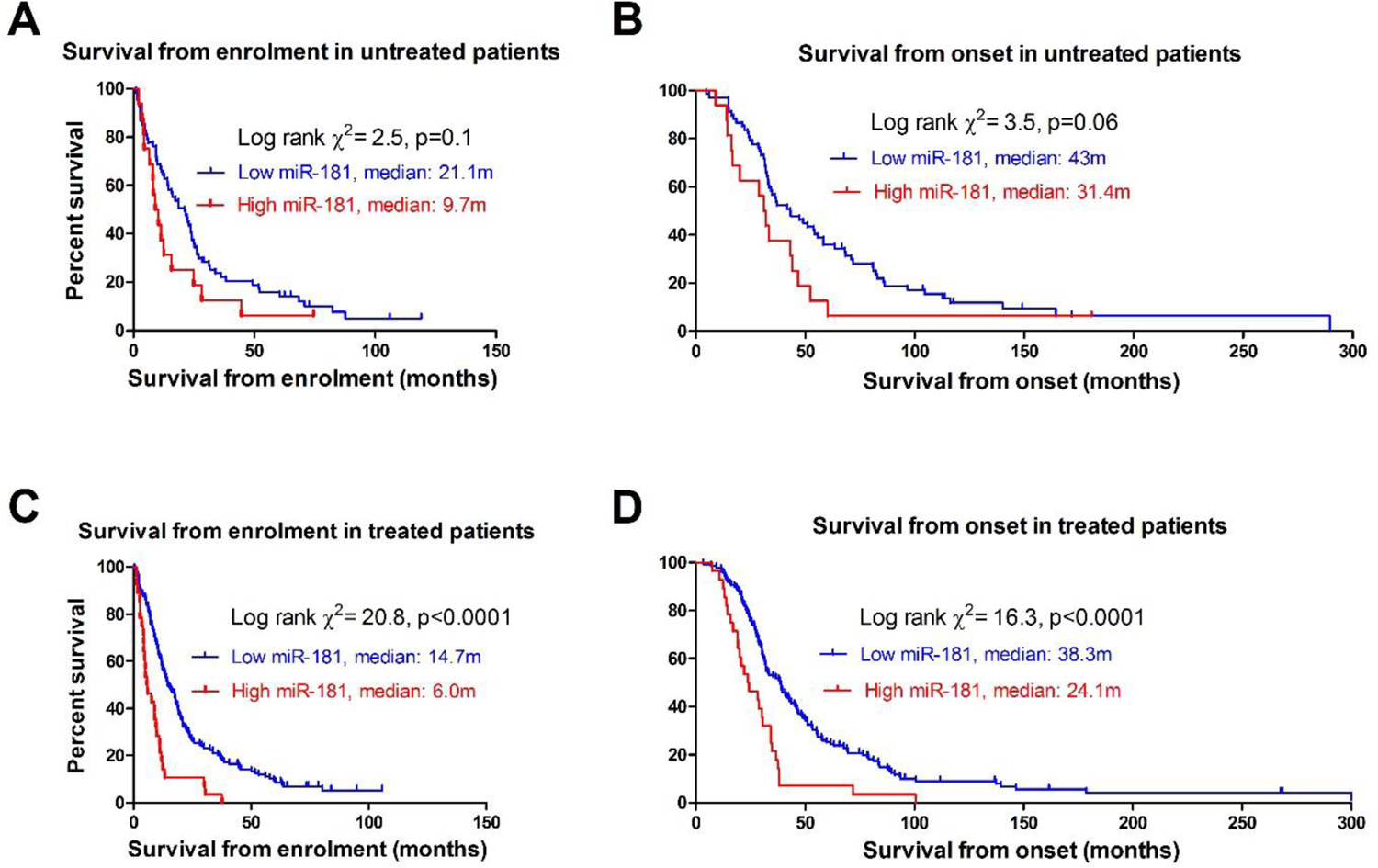
miR-181 levels are predictive of survival length in both untreated patients, from enrolment **(A)** or onset **(B)**, and in treated patients, from enrolment **(C)** or onset **(D).**

**Figure S8.**
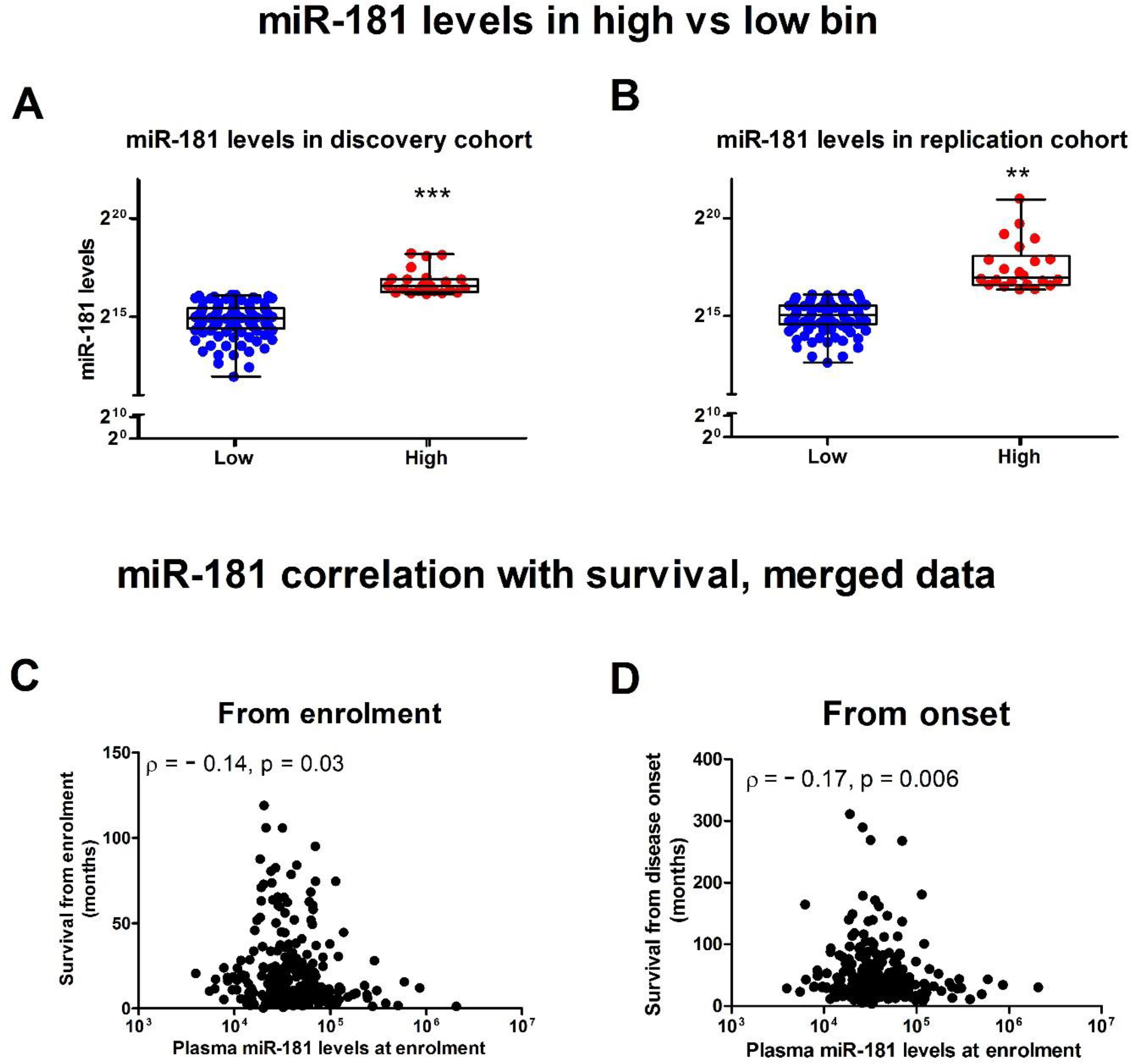
miR-181 levels in the high vs low expression bin, in the **(A)** discovery cohort and **(B)** replication cohort. Plots depicting inverse correlation between miR-181 levels and survival from first phlebotomy **(C)**, or from disease onset **(D)**. **p<0.01, ***p<0.001, t-test with Welch’s correction.

**Figure S9.**
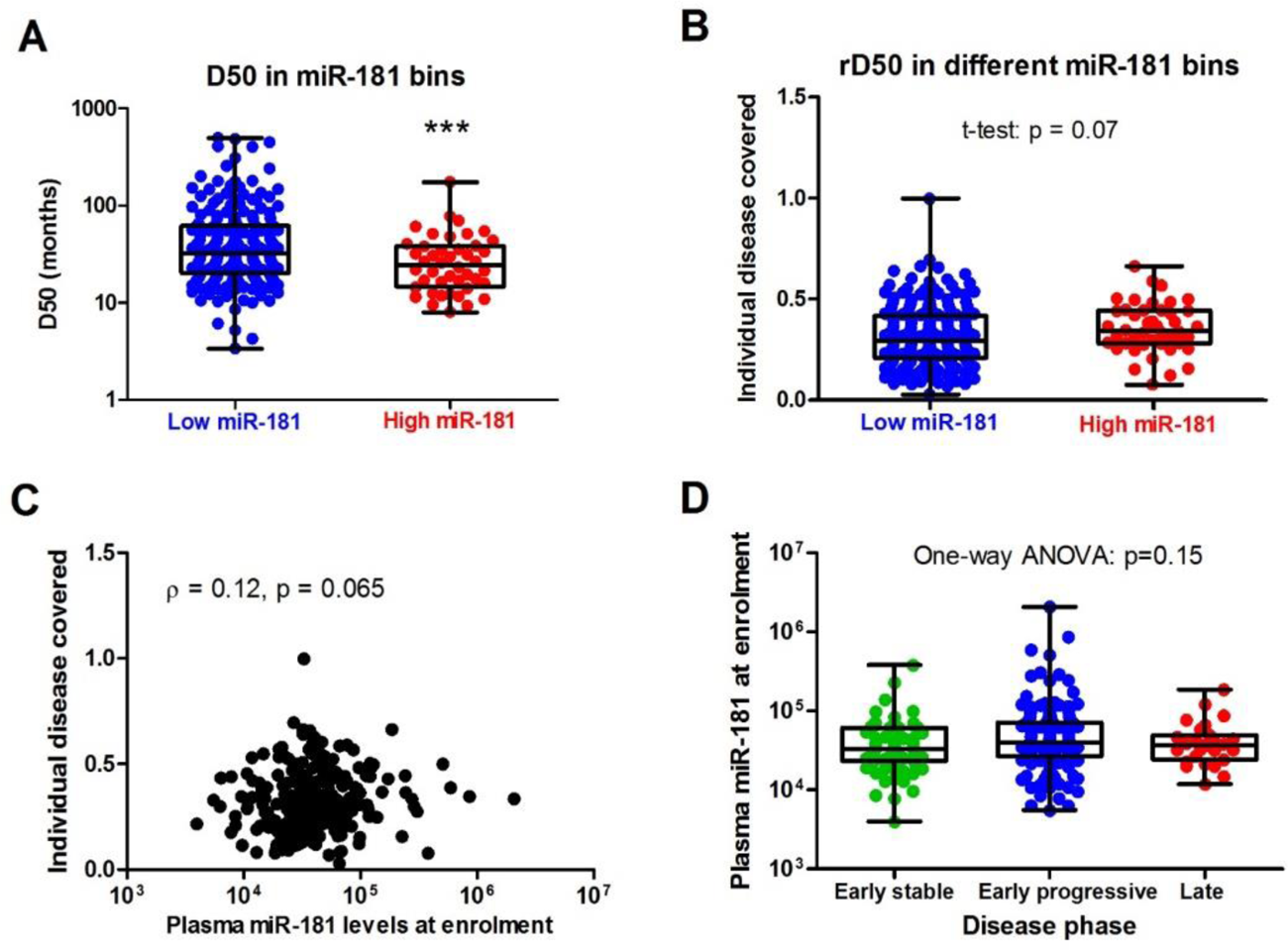
miR-181 levels with respect to parameters of the D50 model. **(A)** D50, a measure of disease aggressiveness, is significantly lower in high vs low miR-181 levels, indicating a more aggressive disease. **(B)** Individual disease covered, reflected by rD50 values, is not different between low and high miR-181 expression bins. **(C)** No correlation of miR-181 levels with individual disease covered. **(D)** miR-181 levels are not different between different phases of disease defined by rD50 values. ***p<0.001, t-test with Welch’s correction.

**Figure S10.**
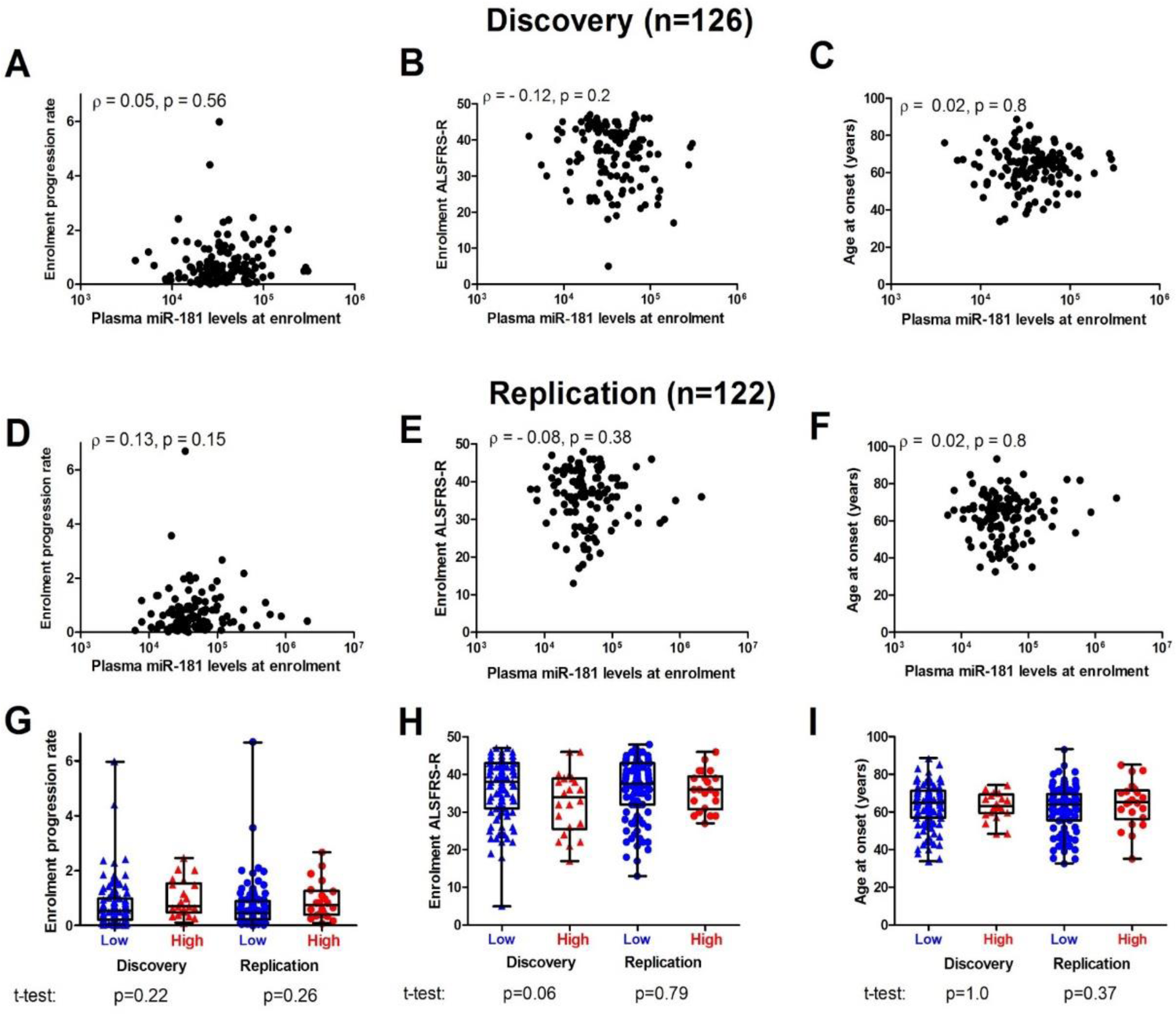
miR-181 levels at enrolment are not correlated with phenotypic properties in the discovery cohort **(A-C)**, or in the replication cohort **(D-F)**. These properties were not different between low and high miR-181 bins **(G-I)**.

**Figure S11.**
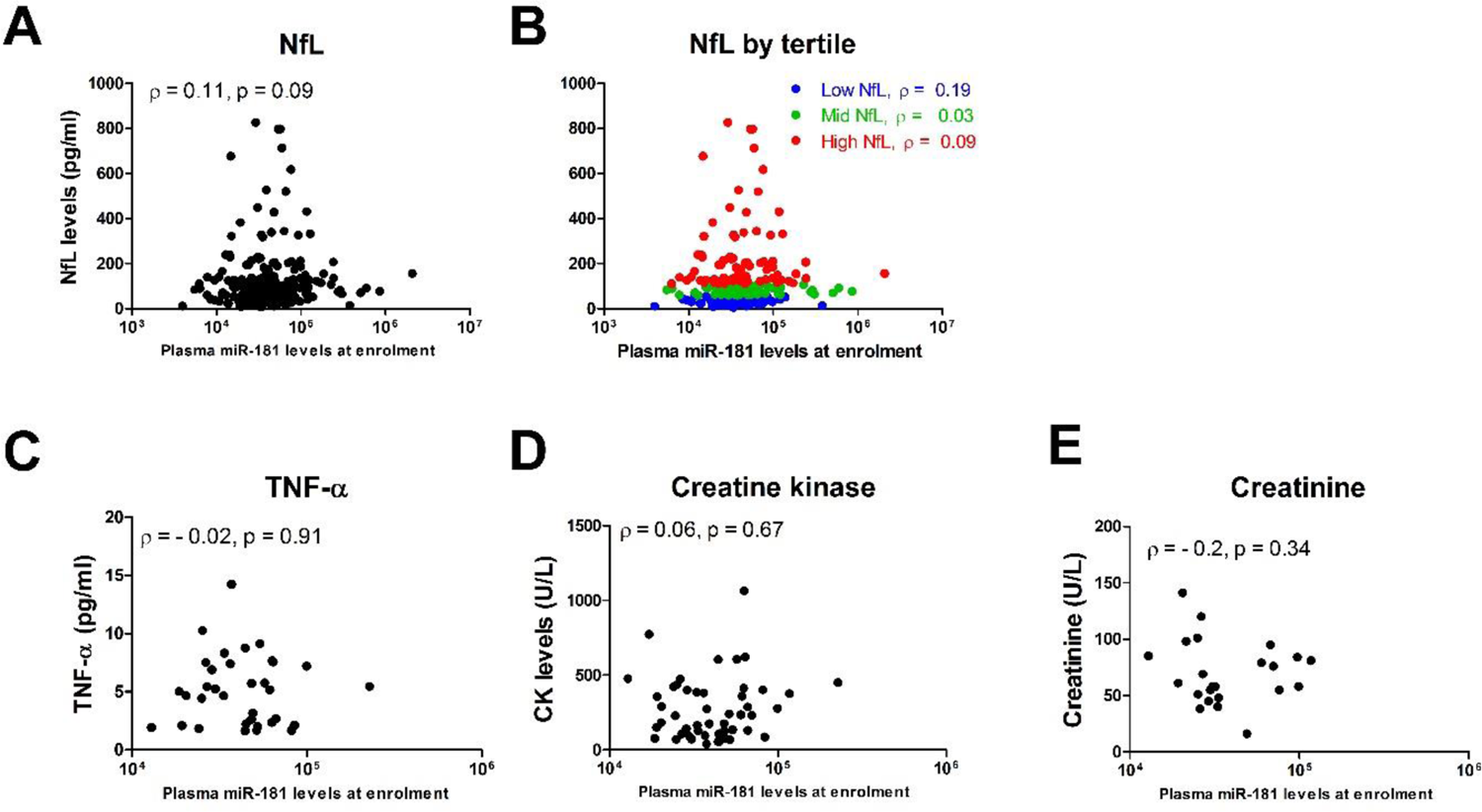
Association of miR-181 with other markers. miR-181 is not correlated with NfL levels, either in the full cohort **(A)** or when NfL is broken into tertiles **(B)**. **(C-E)** miR-181 is not correlated with markers of muscle integrity (CK and creatinine) or inflammatory marker (TNF-alpha).

## Notes

### Competing Interest Statement

The authors have declared no competing interest.

